# Comprehensive benchmarking of metagenomic classification tools for long-read sequencing data

**DOI:** 10.1101/2020.11.25.397729

**Authors:** Josip Marić, Krešimir Križanović, Sylvain Riondet, Niranjan Nagarajan, Mile Šikić

**Author notes:** Authors contributed equally (Josip Marić, Krešimir Križanović).

## Abstract

**Background:** Long reads have gained popularity in the analysis of metagenomics data. Therefore, we comprehensively assessed metagenomics classification tools on the species taxonomic level. We analysed kmer-based tools, mapping-based tools and two general-purpose long reads mappers. We evaluated more than 20 pipelines which use either nucleotide or protein databases and selected 13 for an extensive benchmark. We prepared seven synthetic datasets to test various scenarios, including the presence of a host, unknown species and related species. Moreover, we used available sequencing data from three well-defined mock communities, including a dataset with abundance varying from 0.0001% to 20% and six real gut microbiomes.

**Results:** General-purpose mappers Minimap2 and Ram achieved similar or better accuracy on most testing metrics than best-performing classification tools. They were up to ten times slower than the fastest kmer-based tools requiring up to four times less RAM. All tested tools were prone to report organisms not present in datasets, except CLARK-S, and they underperformed in the case of the high presence of the host’s genetic material. Tools which use a protein database performed worse than those based on a nucleotide database. Longer read lengths made classification easier, but due to the difference in read length distributions among species, the usage of only the longest reads reduced the accuracy.

The comparison of real gut microbiome datasets shows a similar abundance profiles for the same type of tools but discordance in the number of reported organisms and abundances between types. Most assessments showed the influence of database completeness on the reports.

**Conclusion:** The findings indicate that kmer-based tools are well-suited for rapid analysis of long reads data. However, when heightened accuracy is essential, off-the-shelf mappers demonstrate slightly superior performance, albeit at a considerably slower pace. Nevertheless, a combination of diverse categories of tools and databases will likely be necessary to analyse complex samples. Discrepancies observed among tools when applied to real gut datasets, as well as a reduced performance in cases where unknown species or a significant proportion of the host genome is present in the sample, highlight the need for continuous improvement of existing tools. Additionally, regular updates and curation of databases are important to ensure their effectiveness.

## Background

The main task in metagenomic sample analysis is determining its composition – organisms present in the sample and their quantity. The accuracy of the final result depends on many factors, including contamination with other genetic material (i.e. host’s DNA), material isolation, sequencing preparation, and used sequencing technology and classification tools. The recent improvement in both the length and accuracy of long-read sequencing technologies promises a more precise analysis. The advent of high-throughput sequencing has enabled a detailed analysis of microbial communities and their hosts through metagenomics [1,2]. Together with genetic material isolation, an essential component of metagenomics sequencing workflows is a computational method for recognising organisms in a sample. Most current methods are originally designed to work with short, accurate reads from second-generation sequencing technologies. However, due to an increase in accuracy and throughput, long-read sequencing technologies are gaining popularity. Pacific Biosciences (PacBio) and Oxford Nanopore Technologies (ONT) are the most popular long-read sequencing technologies.

Detection of microbes and their abundances using sequencing technologies can be divided into marker gene (typically 16S rRNA) sequencing [3] and whole-metagenome shotgun sequencing. Since the 16S rRNA gene consists of both conserved and variable regions, it is suitable for cost-effective bacteria and archaea profiling.

On the other hand, whole-metagenome shotgun sequencing covers all genomic information in a sample. It enables additional analyses such as binning into Metagenome assembled genomes (MAGs) or previously reconstructed genome sequences, antibiotic resistance gene profiling, and metabolic function profiling. Metagenomic analysis pipelines often begin by detecting and quantifying the taxa in a sample. When most of the genomes present in the sample are unknown, metagenomic de novo assembly methods (i.e. [4]) are the preferred approach. Otherwise, one can compare sequenced data, mapping it to a reference database that stores genomic information related to various taxa in FASTA format.

This work aims to analyse the performance of methods based on comparing long-read sequencing data with a reference database. Several recent studies have proved the value of using long reads in the metagenomic analysis [5–7]. Although there are several benchmarking papers on long reads [7–10], our evaluation additionally includes HiFi PacBio reads, and standard long reads mappers and incorporates an assessment of the influence of the database, read lengths and definition of abundance measures on the results in simulated and real use cases. We also assessed trade-offs between running time and memory requirement.

We evaluated the performance of thirteen different pipelines using seven synthetic datasets, three datasets obtained from mock communities and six real gut microbiome datasets. Some tools are based on nucleotide databases, while others need protein databases.

## Results

Tested metagenomics classification pipelines could be roughly divided into four groups:

(1) kmer-based (Kraken2 [11], Bracken [12], Centrifuge [13], CLARK [14], CLARK-S [15]),
(2) mapping-based (MetaMaps [16], MEGAN-LR with a nucleotide database [17], deSAMBA [18]) tailored for long reads,
(3) general purpose long read mappers (Minimap2 [19] in alignment and mapping mode, Ram [20] in mapping mode),
(4) and protein database-based (Kaiju [21] and MEGAN-LR with a protein database [17]).

To distinguish between two MEGAN-LR versions, the one which uses a protein database and the one which uses a nucleotide database, we named them MEGAN-P and MEGAN-N, respectively. Bracken [12] is a statistical method that computes species abundance using taxonomy labels assigned by Kraken/Kraken2. We adapted Minimap2 and Ram for metagenomics classifications. Mappers usually enable full alignment mode (calculating alignment path) and mapping mode (calculating approximate alignments). We assessed Minimap2 in both modes and Ram only in mapping mode.

We also have reviewed k-SLAM [22], MetaPhlAn [23], ConStrains [24], PathoScope [25], KrakenUniq [26], Sigma [27], CCMetagen [28], and Gotcha [29], but they either crashed during the database creation or performed poorly on long-read data. MEGAN-LR using the DIAMOND [30] aligner and a protein database achieved worse results than MEGAN-LR with the LAST aligner (MEGAN-P), so we omitted it from the analysis. We did not evaluate web-based tools such as BugSeq [31].

There are two main requirements for classification algorithms: identifying present organisms using a taxonomic rank (species in this work) and evaluating their abundances. Reaching these objectives highly depends on the community’s content and the number of reads for each organism. Therefore, using existing PacBio and ONT reads, we synthesised several simple to more complex communities containing 3 to 50 species, varying from highly abundant to very sparse.

– Datasets ONT1, PB1, and PB4 reflect a community of bacteria without eukaryotic species.
– Datasets ONT2 and PB2 reflect metagenomics datasets with one or more eukaryotic species and many bacterial species, respectively.
– Dataset PB3 reflects a community with predominantly human reads (99 %) and two low-abundance bacterial species, reflecting what one might see in an infection setting.
– Datasets PB1+NEG and PB2 represent a situation where a significant portion of the reads comes from an organism that is not present in the database and has no similar present organisms. For the PB1+NEG dataset, we created randomized reads from the human genome. For the PB2 dataset, we added reads from *D. melanogaster* and human isolate datasets.

We also used three well-defined mock community datasets PB_Zymo (PacBio HiFi sequenced Zymo Gut Microbiome Standard dataset), ONT_Zymo (ONT sequenced Zymo Gut Microbiome Standard dataset) and PB_ATCC (PacBio HiFi sequenced ATCC sample). It is important to notice that while we used reads sequenced with older PacBio technologies for synthesised communities, mock communities were sequenced using Sequel 2 HiFi technology. In all tests with mock community datasets, we included PB_Zymo, although the differences between reported and expected values were much larger than for the other two datasets. Due to this disproportion, we will focus more on other datasets in our analysis and conclusion.

Finally, we compared the tools using six real gut datasets. Three datasets are sequenced by ONT technologies (SRR15489009, SRR15489011 and SRR15489017) and three by PacBio HiFi (Sample10, Sample20 and Sample21).

The tools were tested in four areas:

1. Read level classification – how accurately can they classify each read.
2. Abundance estimation – how well can they be used to estimate the abundance of organisms in the sample.
3. Organism detection – how accurately can they detect organisms in a sample.
4. Computational resource usage – running time and consumption of RAM.

We focus our analysis on microbial species in all areas mentioned above apart from read-level classification, where we assessed tools on species and genus ranks. Accuracy and abundance errors were calculated only for the microbial organisms, ignoring reads assigned to the human.

### Read-level classification

We assessed the tools’ read-level classification accuracy on seven synthesised datasets. In addition to species taxonomy rank, we analysed read accuracy on genus level to evaluate the performances of mappers when some of the species in the sample were not present in the database. Figure 1 shows that off-the-shelf mappers and mapping-based tools outperformed others on almost all datasets and both levels. Differences between them and kmer-based tools varied up to 10 % at species levels. The only exception was MEGAN-N which performed similarly to kmer-based tools. Pipelines which use a protein database underperformed significantly.

**Figure 1.**
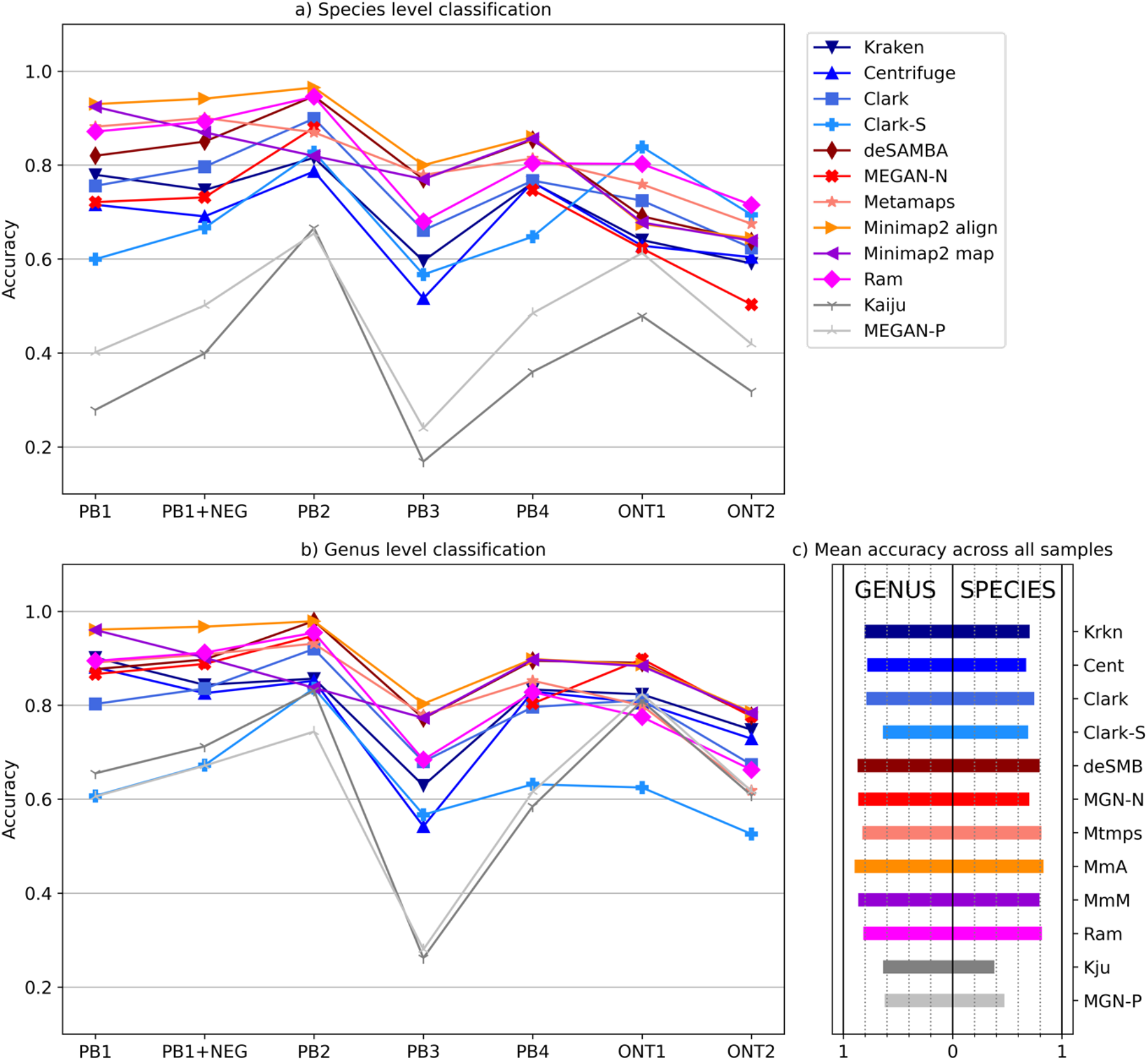
Read level classification accuracy, comparison between species and genus level classification. Kmer-based are represented in red, mapping-based tools are represented in blue and protein tools are represented in green. Plot a) shows species-level classification for which reads are considered correctly classified if classified to a correct species. Plot b) shows genus-level classification for which reads are considered correctly classified if classified to a correct genus. Plot c) shows mean values for both levels. Results for MEGAN-N are unavailable for the PB3 dataset.

Minimap2 with alignment outperformed other tools, followed by mapping-based tools (deSAMBA, Metamaps) and Ram. Interesting cases were ONT1 and ONT2 datasets which contain reads of two species of the Vibrio genus that were not in databases. Since there were other similar species of the Vibrio genus in the database, some tools, such as MEGAN-N and Minimap2 in both modes, tended to assign those reads to them. In contrast, other tools, such as CLARK-S and Ram, tended to leave those reads unassigned. Therefore, the results on the ONT1 and ONT2 datasets for these four tools were almost reversed when analysing genus and species levels. CLARK-S and Ram had the highest accuracy when inspecting the ONT1 and ONT2 datasets at the species level and among the lowest when examining the dataset at the genus level. In contrast, Minimap2 and MEGAN-N had the highest accuracy at the genus level but performed worse at the species level.

Since there is an imbalance in the number of reads per species, we additionally calculated the F1 score for each class (organism in the sample) separately and averaged them (F1 macro average). Using F1 macro average instead of accuracy shows a similar pattern for most datasets with Minimap2 (both modes) surpassing others and a narrower distance between mapping-based and kmer-based tools (Supplementary Figure 1).

We further investigate the influence of read length on classification. We only present analysis for Minimap2 with alignment, the most accurate tool at the read level. As evident from Figure 2, increasing the read length led to a higher classification level. However, we could not select only the longest reads due to different read length distributions per organism. Additional analysis on how using 30% of longest reads impacted the results is provided in Supplementary Table 1.

**Figure 2.**
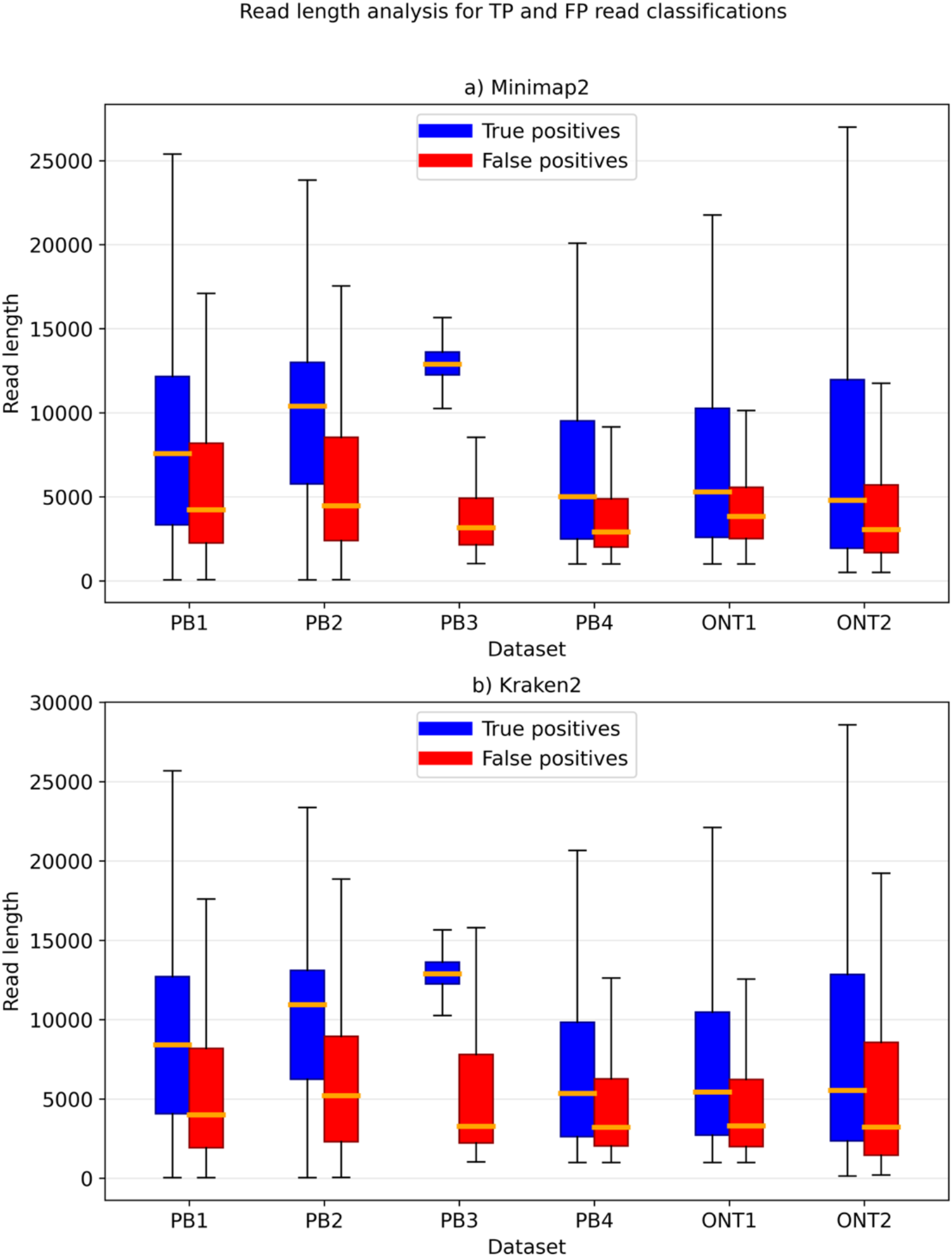
Comparison between classification accuracy and read length. The figure shows Q1, median and Q3 read length for true positive and false positive read classifications for each dataset. False positive read lengths are considered only for organisms in the database. The results shown in the figure were obtained using Minimap2 with alignment and Kraken.

### Abundance estimation

Abundance estimation is arguably the most important assessment. Tools report different types of abundance. Most assessed tools report relative read counts as abundance. However, abundance might be calculated in different ways. Other measures count genomes or cells. The difference between these two is that the latter takes into account ploidy. The vendors of mock community samples usually report all of them. Here we also tested a method that predicts genomic DNA length using read lengths. This is irrelevant for short-read sequencing because all reads are the same size. However, long-read sequencing technologies produce reads whose length might vary greatly, especially for ONT technology.

In the analysis, we compared L1 distances between reported and declared abundances using relative read count, genome length and genome count. Supplementary Table 5 shows the theoretical composition of Zymo and ATCC datasets.

Table 1 shows that the results are mostly consistent, considering different ways of abundance calculation. The ranking of tools by different measures slightly varies. However, it is important to note the difference between read counts and genome length measures. Although distances do not vary much for PacBio HiFi data and estimations of genomic DNA abundances are slightly more accurate when read counts are used, differences for ONT reads are noticeable and adding the length of reads increases the accuracy of the prediction. The reason for this behaviour might be in read distributions. While it is narrow for PacBio HiFi reads, it is wide and skewed for ONT reads.

**Table 1.**
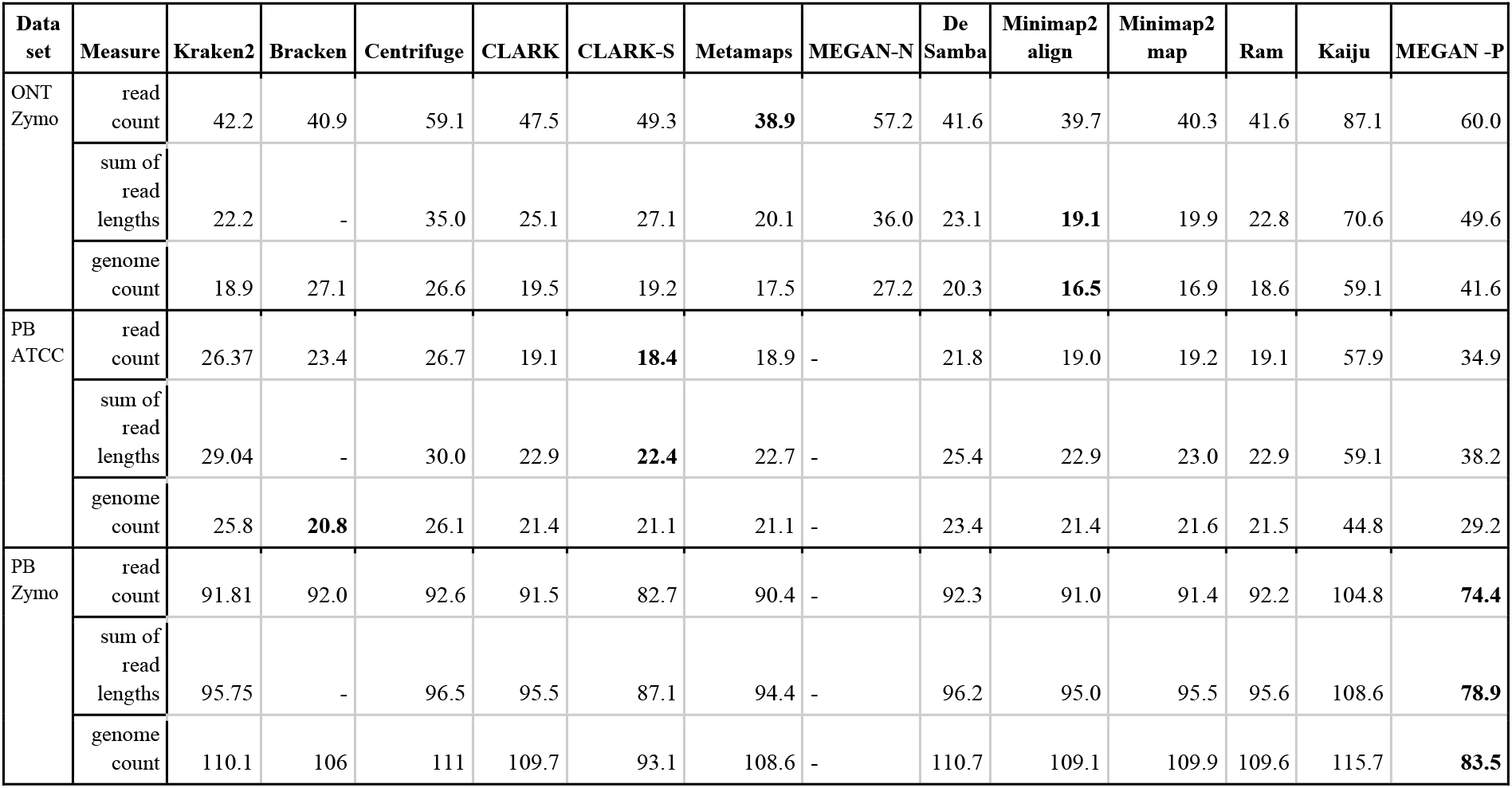
Comparison of the total abundance estimation error between relative read count, relative genome length and relative genome count abundances. The table shows the total abundance estimation error for mock community datasets for three definitions of relative abundance: 1) the percentage of reads classified to a species, 2) the percentage of genome lengths (sum of read lengths) classified to a species and 3) the percentage of genomes classified to a species. The total error was calculated as the L1 distance between specific types of abundances reported by tools and the abundances declared by vendors and summed up across all organisms (all present and all reported). The lowest abundance errors are in bold. Unfortunately, due to its long-running time, results for MEGAN-N are unavailable for datasets PB_Zymo and PB_ATCC. It is important to note that Bracken produces only read counts assigned to a taxonomic rank. Therefore, to compare it with other tools, abundances – the percentage of genomes of species in the sample, were calculated by normalising read counts with the average genome length of the species to which corresponding read counts were assigned.

Inspired by medical laboratory tests which use cell count to measure the abundance, we continue with relative genome count measure in the rest of the paper.

In the next test, we evaluated the tools on databases with and without the human genome (containing only bacterial and archaeal genomes) for datasets with a higher percentage of the human genome. The comparison of the abundance estimation error is given in Table 2.

**Table 2.** Comparing abundance estimation error for the database with the human genome and the database without the human genome. The table shows the total abundance estimation error for datasets PB2, PB3 and ONT2 (datasets that contain human reads) for all tools and two databases: a database with the human genome and a database without the human genome. The error was calculated as the L1 distance between abundances calculated for each tool and true abundances. Each dataset name is followed by the percentage of human reads in that dataset in parentheses.

| Dataset (% human) | Database | Kraken2 | Bracken | Centrifuge | CLARK | CLARK-S | deSAMBA | Metamaps | MEGAN-N | Minimap2 align | Minimap2 map | Ram | Kaiju | MEGAN -P |
| --- | --- | --- | --- | --- | --- | --- | --- | --- | --- | --- | --- | --- | --- | --- |
| ONT2 (5.78%) | human | 79.8 | 88.3 | 84.0 | 74.5 | 74.7 | 70.8 | **64.6** | 81.8 | 68.2 | 72.5 | 68.7 | 113.3 | 101.5 |
|  | no human | 83.6 | 88.1 | 90.6 | 75.9 | 74.7 | 74.9 | **64.6** | 84.2 | 72.3 | 76.0 | 69.6 | 113.5 | 101.4 |
| PB2 (20%) | human | 43.7 | 22.5 | 61.5 | 16.3 | 23.2 | 14.1 | 25.8 | 17.6 | 7.8 | 81.4 | **6.4** | 51.5 | 73.9 |
|  | no human | 89.2 | 54.2 | 98.3 | 63.8 | 23.3 | 68.8 | 25.9 | 38.9 | 48.9 | 106.8 | **8.8** | 57.3 | 78.1 |
| PB3 (99%) | human | 50.1 | 21.3 | 104.8 | 35.5 | **20.8** | 32.0 | 23.7 | 63.7 | 27.8 | 117.7 | 61.6 | 139.5 | 144.6 |
|  | no human | 145.7 | 145.1 | 145.7 | 145.7 | 140.1 | 145.7 | **96.0** | 145.7 | 145.5 | 145.5 | 140.3 | 141.4 | 145.1 |

Table 2 shows that when the human reads’ percentage is lower, tools such as CLARK-S, Ram, Metamaps, Kaiju, and MEGAN-P were unaffected by the presence of the human genome in the database. However, as the human genome reads’ percentage approaches 100 %, the accuracy of most tools significantly declines. Exceptions were tools which used protein databases, and their abundance estimation slightly changed. In the case of the highly represented human genome, Metamaps achieved notably better results than other tools. However, its error almost quadrupled. Therefore it is important for each experiment to prepare a database which would include all organisms (including the host) that might be contained in the sample.

There was no clear winner when we included the human genome in databases. Metamaps performed best on ONT2, Ram on PB2, and CLARK-S on PB4. Centrifuge and protein-based tools were significantly worse than others. It is important to note that Bracken significantly improved Kraken2 results for PacBio datasets.

To analyse abundance error in more detail, we separately calculated the error for species present in the sample and for species not present in the sample but incorrectly reported by tools. We calculated the L1 distance between reported and expected abundance in percentages. Figure 3 and Supplementary Figure 2 show the results.

**Figure 3.**
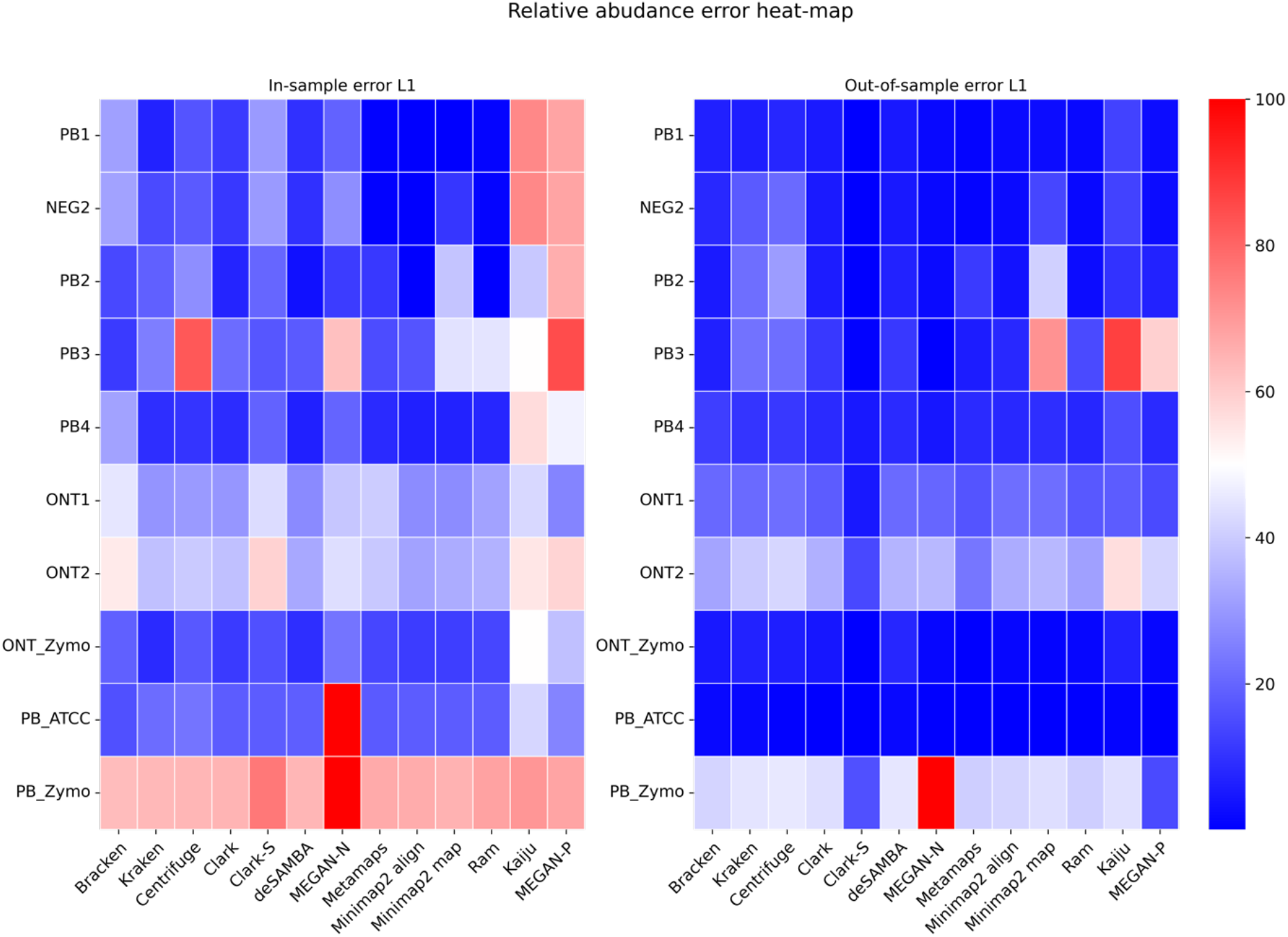
Abundance estimation error on species level – heatmap. Abundance estimation error was calculated by comparing the abundances calculated for each tool to the ground truth. Errors were calculated separately within the dataset and outside the dataset. The total abundance estimation error is represented by the colour intensity of the corresponding cell. The red colour represents a higher error, and the blue represents a lower error. It is important to note that in the case of the PB_Zymo dataset, species *Veillonella rogosae* (taxId: 423477) and *Prevotella corporis* (taxId: 28128), which represent 19.94 % and 6.26 %, respectively, were not present in the database.

Minimap2 align outperformed other tools in absolute differences between abundances of present organisms. In most of the datasets, its mean difference was below 2%, yet, other tools were not far away. Tools did not achieve good results on the PB_Zymo dataset. One of the reasons is the lack of two species in databases, *Veillonella rogosae* and *Prevotella corporis,* which represent more than 26 % of the dataset. However, even if we did not consider these two species, distributions of errors for this dataset were much wider than in another PacBio mock community dataset, PB_ATCC (Supplementary Figure 2).

Regarding species not present in the dataset, CLARK-S surpassed others, followed by MetaMaps, Ram and MEGAN-N. Minimap2 was more prone to reporting organisms not present in the sample. Therefore we deem there is space for improvement in the postprocessing analysis (e.g. using sequence similarity threshold or EM approach similar to MetaMaps) or by changing parameters such as kmer length or the percentage of filtered kmers.

### Organism detection

In the analysis of microbial communities, it is often important to accurately detect present microbes. Table 3 shows how the number of true and false positive organism detections is related to a threshold – a minimal number of assigned reads required to consider an organism as detected. The table shows that kmer-based tools, Kraken2 and Centrifuge, often reported a huge number of species, usually an order of magnitude more than mapping-based tools and mappers, except the Minimap2 map. CLARK-S surpassed other tools for all datasets, followed by Ram and Bracken. Since we did not have the same mock community sequenced with the same coverage with ONT and PacBio HiFi reads, comparing sequencing technologies was difficult. What is evident is that kmer and protein-based tools reported a significantly lower number of organisms for mock communities sequenced by PacBio HiFi than with ONT. In comparison, mappers and mapping-based tools reported a similar or slightly lower number of organisms. As expected, increasing the threshold for most datasets decreases the number of false positive detections while retaining the same number of true positive detections. In the case of very low abundance species, such as in datasets PB4 (lowest proportion of reads – 0.005 %) and ONT1 (lowest proportion of reads – 0.01 %), PB_Zymo (lowest proportion of reads – 0.0001%) and PB_ATCC (lowest proportion of reads – 0.02%) increasing the threshold lowered the number of true positive detections.

**Table 3.**
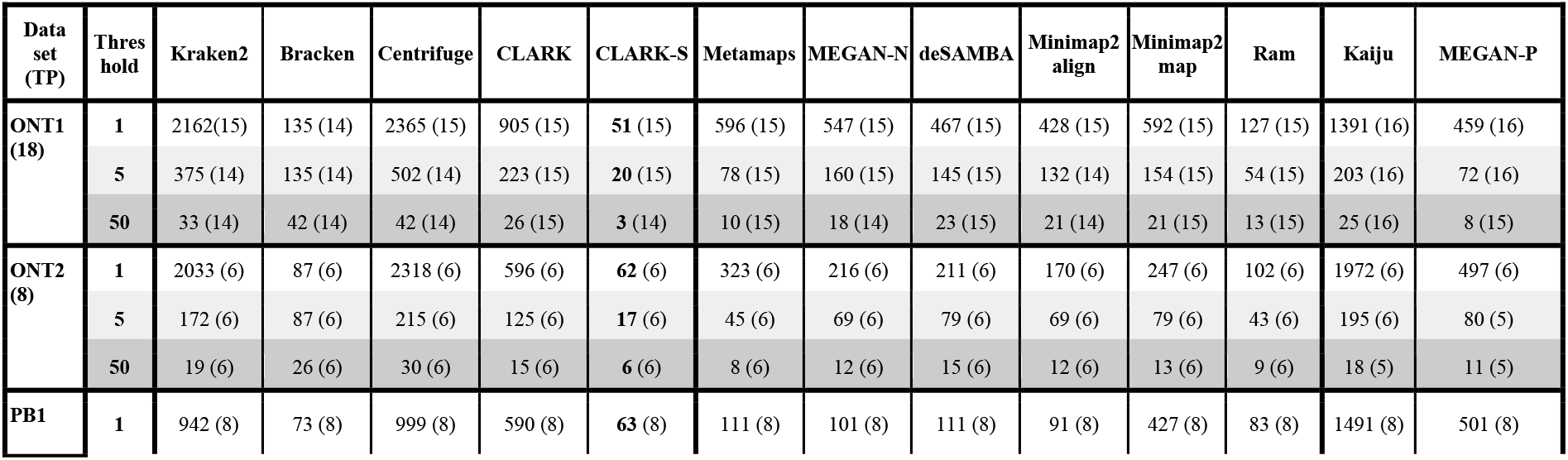
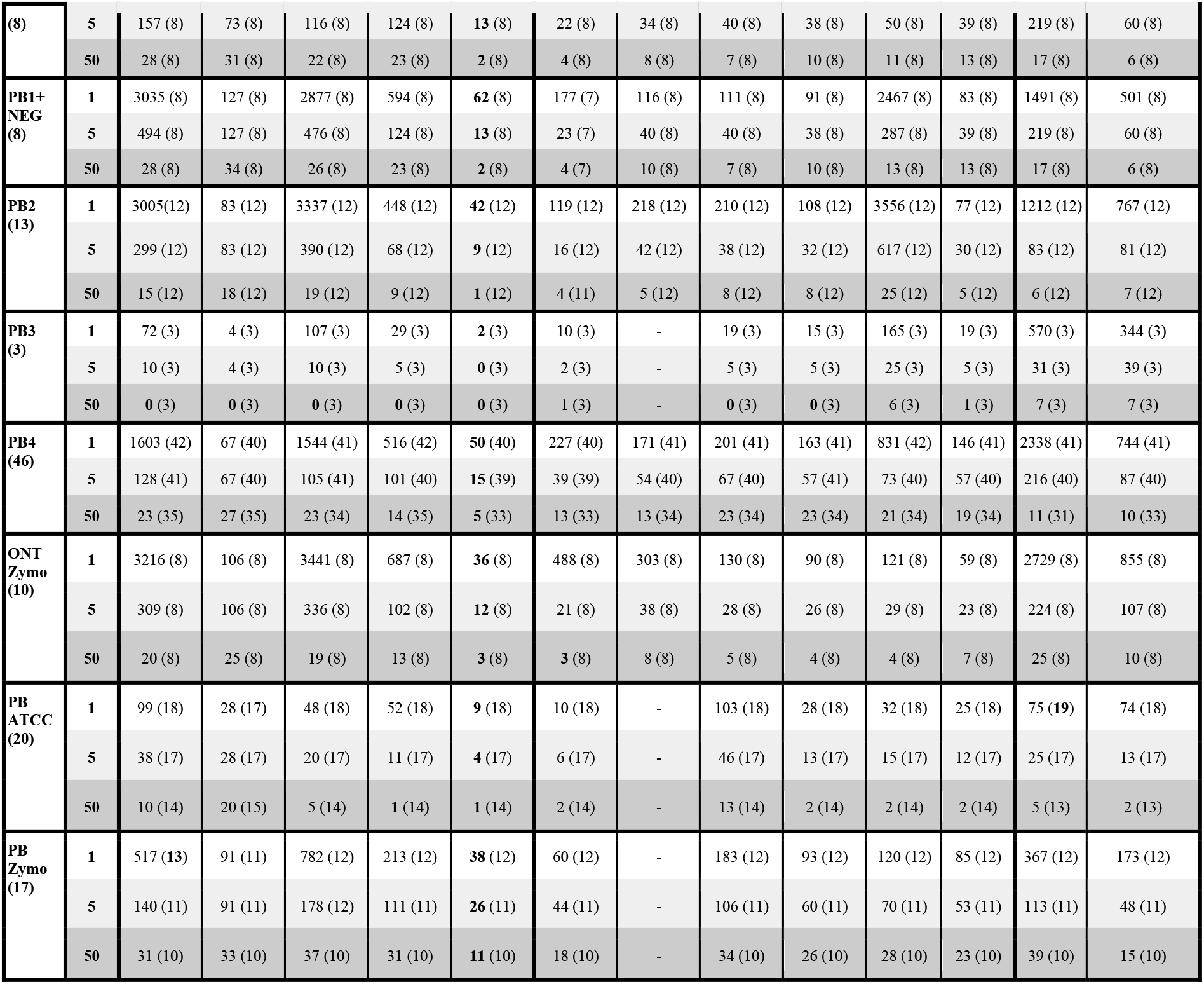
True positive and false positive organism detection. The table shows true and false positive organism detections for three different thresholds: 1, 10 and 50. A threshold represents the number of reads that need to be assigned to that organism to consider it present in the sample. The data is presented as the number of false-positive detections (organisms incorrectly reported as present), followed by the number of true positive detections in parentheses (organisms correctly reported as present). Each dataset name is followed by the number of species in that dataset in parentheses. Results for MEGAN-N are unavailable for datasets PB3, PB_Zyme and PB_ATCC.

Additionally, we analysed abundances and the number of correctly identified organisms for those species with abundances lower than 1 %. Results, presented in Supplementary Table 9, show that there is no clear winner in the abundance estimation accuracy. Yet, Minimap2 align outperformed others in two out of four datasets. Furthermore, Kraken2 and Minimap2 recognised the most present organisms in all samples except PB_Zymo, where Kraken2 correctly predicted one more. Unfortunately, for mock communities, we did not have information about incorrectly classified reads which belong to known species with a low abundance. Therefore we could not calculate the number of incorrectly classified organisms for the subsets of low-abundant species.

### Real data without ground truth

We also compared pipelines on six real gut microbiome datasets, for which the ground truth was unknown. We compared tools in two ways, with the results viewed as vectors. Firstly, a distance matrix between the tool results was calculated, and a hierarchical clustering algorithm was applied. Secondly, a PCA algorithm was performed on the tools’ outputs, obtaining the two most significant components. The results of both analyses for datasets Sample10, Sample20, Sample21, SRR15489009, SRR15489011 and SRR15489017 are shown in Figure 4.

**Figure 4.**
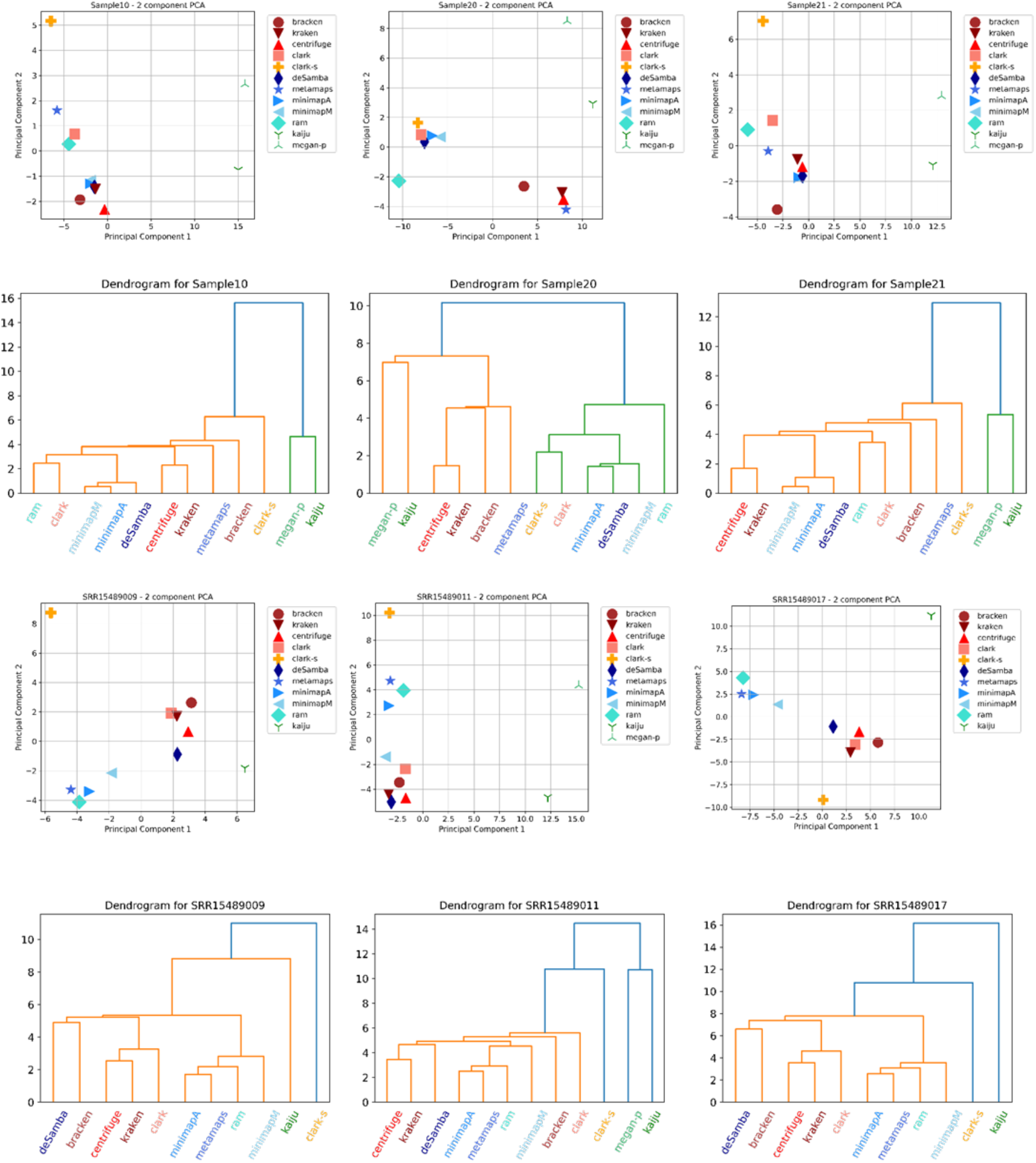
2D PCA and dendrograms for real datasets. Abundance estimation data for each tool is viewed as a vector, with components being abundance estimations for each organism (in and out of the sample). The data was transformed using PCA, and the two most significant components were plotted. Hierarchical clustering was also performed on initial vectors. The Figure displays 2D PCA plots and hierarchical clustering dendrograms for all real datasets. Data for MEGAN-N is unavailable on all datasets due to dataset sizes. Data for MEGAN-P is unavailable on datasets SRR15489009 and SRR15489017.

From Figure 4, it can be seen that kmer-based tools such as Kraken2, Bracken and Centrifuge performed similarly. Clark was very close for most samples. While for PacBio HiFi datasets, mappers Minimap2 and Ram, and Metamaps performed similarly, for ONT datasets Ram and Metamaps performed differently than Minimap2 tools. deSAMBA was usually somewhere in between these kmer-based and mapping-based tools but usually closer to kmer-based tools. CLARK-S, Kaiju and MEGAN-P were far from other tools.

We analysed the ten most abundant species for each pipeline for the SRR15489009 (PacBio) dataset to better understand the differences between pipelines. Results are presented in Supplementary Table 11. Here, we saw a significant difference in content between protein and nucleotide databases. Although belonging to the core [32] of the human gut, *Lachnospiraceae bacterium* species are not found in the protein database. It is important to note that CLARK and CLARK-S did not recognise it, although they use the nucleotide database, which contains them. Protein database does not contain another prevalent [33] human gut bacteria – *Eubacterium rectale*. Furthermore, while mappers and all mapping-based tools recognised it (average abundance of 5 %), none of the kmer-based tools did the same. Other important human gut species, such as *Prevottela copri* [34] and *Ruminococcus torques* [35], were not present in the NCBI-NT database.

During the analysis of the SRR15489009 (PacBio) dataset’s results, we noticed significant discordance in abundances for *Faecalibacterium prausnitzii* [36] between tools. While kmer-based tools’ abundances were 12%-14% (CLARK-S with 25% is an outlier), most mappers and mapping base tools’ abundances were 17%-18%. Kaiju and MEGAN-P report 11% and 19%, respectively.

A further interesting question for real gut datasets is the actual number of species present in the sample. This depends on the coverage. However, the total read length for PacBio HiFi and ONT datasets is similar, so we compared the reported results. Supplementary Table 3 shows the percentage of reads classified for all datasets and tools. The least number of classified reads were for three ONT gut samples, where best-performing tools rarely classified above 50% of reads. The numbers for PacBio samples were even above 80% for some tools. This might be because of the higher error rate and shorter read length of ONT data. Although there is a difference between the PacBio HiFi datasets (PB_Zymo and PB_ATCC), it is much smaller. The remaining 20 % of unclassified reads might be explained by missing species in databases (Supplementary Table 10).

While Supplementary Table 4 shows the number of detected species for each real dataset and each tool, Table 4 shows a more detailed analysis of real gut samples. CLARK-S, Metamaps and Ram reported a significantly lower number of sequences for real gut datasets sequenced by PacBio HiFi technologies than all other tools. Since they performed better for real mock community datasets PB_Zymo and PB_ATCC, we hypothesise the number of species possible to detect for selected sequencing depth was closer to the numbers they reported and probably even smaller.

**Table 4.** The number of detected species for real gut microbiome datasets. The table shows the number of detected species on real datasets for three different thresholds: 1, 10, and 50. A threshold represents the minimum number of reads assigned to a species to consider it present in the sample. Reads assigned to higher taxa are considered unclassified. The table also shows the percentage of unclassified reads for each dataset. Due to dataset sizes, results for MEGAN-N are unavailable. Results for MEGAN-P are unavailable on datasets SRR15489009 and SRR15489017.

| Data set |  |  | Kraken2 | Bracken | Centrifuge | CLARK | CLARK-S | Metamaps | deSAMBA | Minimap2 align | Minimap2 map | Ram | Kaiju | MEGAN -P |
| --- | --- | --- | --- | --- | --- | --- | --- | --- | --- | --- | --- | --- | --- | --- |
| Sample 10 (ONT) | % unclassified |  | 55.5 | 47.3 | 51.8 | 65.3 | 80.2 | 66.0 | 51.4 | 50.8 | 50.6 | 72.1 | 74.2 | 83.8 |
|  | Detected with thres hold | 1 | 5487 | 4775 | 5538 | 2542 | 418 | 4973 | 1017 | 1051 | 1504 | 432 | 7102 | 3046 |
|  |  | 5 | 5160 | 4775 | 5286 | 824 | 171 | 3149 | 407 | 385 | 437 | 276 | 5247 | 762 |
|  |  | 50 | 1060 | 1253 | 1271 | 169 | 89 | 377 | 192 | 183 | 191 | 137 | 418 | 169 |
| Sample 20 (ONT) | % unclassified |  | 71.6 | 66.2 | 68.6 | 82.9 | 85.7 | 87.8 | 76.4 | 74.7 | 74.6 | 91.3 | 89.0 | 95.4 |
|  | Detected with thres hold | 1 | 5459 | 4560 | 5523 | 1736 | 452 | 4562 | 659 | 786 | 996 | 316 | 6868 | 2801 |
|  |  | 5 | 5064 | 4560 | 5225 | 337 | 165 | 2600 | 296 | 298 | 326 | 180 | 4296 | 587 |
|  |  | 50 | 1375 | 1569 | 1596 | 100 | 74 | 478 | 130 | 139 | 140 | 77 | 245 | 136 |

| Data set |  |  | Kraken2 | Bracken | Centrifuge | CLARK | CLARK-S | Metamaps | deSAMBA | Minimap2 align | Minimap2 map | Ram | Kaiju | MEGAN-P |
| --- | --- | --- | --- | --- | --- | --- | --- | --- | --- | --- | --- | --- | --- | --- |
| Sample 10 (ONT) | % unclassified |  | 55.5 | 47.3 | 51.8 | 65.3 | 80.2 | 66.0 | 51.4 | 50.8 | 50.6 | 72.1 | 74.2 | 83.8 |
|  | Detected with thres hold | 1 | 5487 | 4775 | 5538 | 2542 | 418 | 4973 | 1017 | 1051 | 1504 | 432 | 7102 | 3046 |
|  |  | 5 | 5160 | 4775 | 5286 | 824 | 171 | 3149 | 407 | 385 | 437 | 276 | 5247 | 762 |
|  |  | 50 | 1060 | 1253 | 1271 | 169 | 89 | 377 | 192 | 183 | 191 | 137 | 418 | 169 |
| Sample 21 (ONT) | % unclassified |  | 49.0 | 40.4 | 45.7 | 57.5 | 71.3 | 64.3 | 42.1 | 40.6 | 40.5 | 67.2 | 71.3 | 81.6 |
|  | Detected with thres hold | 1 | 5440 | 4944 | 5510 | 2220 | 375 | 4679 | 1010 | 999 | 1390 | 409 | 6837 | 3134 |
|  |  | 5 | 5213 | 4944 | 5315 | 959 | 210 | 3426 | 457 | 444 | 554 | 262 | 5546 | 1249 |
|  |  | 50 | 1774 | 2055 | 2064 | 195 | 102 | 589 | 214 | 207 | 217 | 160 | 743 | 242 |
| SRR 15489009 (PacBio) | % unclassified |  | 15.6 | 5.6 | 14.4 | 18.4 | 49.0 | 40.5 | 12.2 | 39.1 | 24.2 | 46.3 | 12.7 | - |
|  | Detected with thres hold | 1 | 4625 | 2011 | 5177 | 3785 | 294 | 415 | 2197 | 311 | 1930 | 306 | 3184 | - |
|  |  | 5 | 3048 | 2011 | 4151 | 2453 | 172 | 194 | 940 | 229 | 792 | 210 | 1349 | - |
|  |  | 50 | 461 | 654 | 740 | 469 | 97 | 116 | 236 | 138 | 210 | 130 | 388 | - |
| SRR 15489011 (PacBio) | % unclassified |  | 17.4 | 4.5 | 15.9 | 23.8 | 63.1 | 55.6 | 15.7 | 52.9 | 33.3 | 60.7 | 12.1 | 58.5 |
|  | Detected with thres hold | 1 | 4599 | 2452 | 5193 | 3768 | 338 | 460 | 2049 | 375 | 1873 | 351 | 3071 | 1145 |
|  |  | 5 | 2964 | 2452 | 4097 | 2372 | 192 | 227 | 895 | 263 | 771 | 249 | 1355 | 450 |
|  |  | 50 | 564 | 723 | 905 | 555 | 111 | 136 | 285 | 159 | 230 | 150 | 399 | 161 |
| SRR 15489017 (PacBio) | % unclassified |  | 18.1 | 4.5 | 17.3 | 20.2 | 56.0 | 44.9 | 13.1 | 42.9 | 26.7 | 51.4 | 13.8 | ^ |
|  | Detected with thres hold | 1 | 4784 | 2078 | 5260 | 3992 | 347 | 453 | 2383 | 352 | 2149 | 345 | 3506 | ^ |
|  |  | 5 | 3461 | 2078 | 4488 | 2701 | 188 | 224 | 1057 | 252 | 966 | 246 | 1648 | ^ |
|  |  | 50 | 606 | 570 | 1002 | 627 | 107 | 128 | 327 | 146 | 263 | 135 | 509 | ^ |

### Computational resource usage

Results for running time and memory usage are presented in Table 5. As expected, kmer-based tools, apart from CLARK-S and Bracken, surpassed others in the running time. For our synthetic test datasets, Centrifuge has the lowest running time for most datasets. Compared to mappers Minimap2 and especially Ram, the difference between the best kmer-based tools and mappers was below one order of magnitude. Among mapping-based tools, deSAMBA was only a few times slower than the fastest kmer best tools, while MetaMaps and MEGAN-N were among the slowest. Kaiju was among the fastest, and MEGAN-P was among the slowest tools. Ram used the least amount of memory. Kraken2, Centrifuge, Minimap2, MEGAN-P and MEGAN-N, for most datasets, used 2-3 times more memory. deSAMBA, CLARK, CLARK-S and MetaMaps used 10-15 times more memory.

**Table 5.** Resource usage. The table shows running time (in seconds) and memory usage (in GB) for all tools and datasets. The table shows the minimum and maximum values for both measures for all synthetic and real datasets, which are about 1Gbp in size.

|  | Kraken2 | Bracken | Centrifuge | CLARK | CLARK-S | Metamaps | MEGAN-N | deSAMBA | Minimap2 align | Minimap2 map | Ram | Kaiju | MEGAN -P |
| --- | --- | --- | --- | --- | --- | --- | --- | --- | --- | --- | --- | --- | --- |
| **Execution time / s** |  |  |  |  |  |  |  |  |  |  |  |  |  |
| **Min** | 312 | 68971 | 267 | 953 | 3862 | 37839 | 58238 | 895 | 1904 | 1541 | 1272 | 329 | 16183 |
| **Max** | 327 | 224048 | 329 | 993 | 4114 | 145416 | 160225 | 1400 | 4220 | 2283 | 1697 | 610 | 33430 |
| **Memory / GB** |  |  |  |  |  |  |  |  |  |  |  |  |  |
| **Min** | 42.98 | 24.37 | 35.94 | 118.91 | 271.14 | 146.29 | 26.87 | 143.55 | 24.02 | 19.77 | 9.07 | 55.28 | 22.73 |
| **Max** | 43.12 | 45.31 | 37.08 | 120.61 | 271.46 | 208.46 | 108.22 | 213.95 | 47.40 | 38.23 | 14.27 | 55.41 | 93.29 |

**Table 6.** Execution time for datasets of different sizes. Tools were run on three mock community datasets (ONT_Zymo, PB_ATCC and PB_Zymo) subsampled to three different sizes to test their scalability. All datasets used in the main part of the paper were subsampled to 1Gbp (file size is about 2GB) to make the testing viable even for slower tools. However, all three datasets were used in their original size, subsampled to half of the original dataset and subsampled to 1Gbp. The table also shows the execution time for real gut microbiome datasets in their full size. Since Bracken, MEGAN-N, MEGAN-P and MetaMaps running times were prohibitively long on larger datasets, we omitted them from the analysis. Additionally, deSAMBA was unable to process some larger datasets completely. The table shows execution time in seconds.

| Dataset | Dataset size in GB | Kraken2 | Centrifuge | CLARK | CLARK-S | deSAMBA | Minimap2 align | Minimap2 map | Ram | Kaiju |
| --- | --- | --- | --- | --- | --- | --- | --- | --- | --- | --- |
| ONT_Zymo | 2 | 315 | 291 | 954 | 3917 | 1400 | 2874 | 1808 | 1697 | 341 |
|  | 14 | 454 | 779 | 1392 | 7202 | 4092 | 17418 | 7642 | 4122 | 2588 |
|  | 27 | 581 | 1351 | 1923 | 10657 | 6722 | 39334 | 13822 | 6979 | 3780 |
| PB_ATCC | 2 | 312 | 284 | 972 | 3942 | 1666 | 3797 | 1890 | 1303 | 329 |
|  | 55 | 934 | 2679 | 2538 | 15396 | - | 51615 | 20998 | 7552 | 2315 |
|  | 111 | 1316 | 5080 | 6883 | 28756 | - | 104001 | 41232 | 14056 | 4381 |
| PB_Zymo | 2 | 314 | 275 | 974 | 3913 | 1379 | 3145 | 1710 | 1272 | 347 |
|  | 17 | 479 | 841 | 1458 | 7675 | 5548 | 12690 | 5235 | 2984 | 909 |
|  | 34 | 638 | 1487 | 1999 | 11767 | 9930 | 22411 | 9385 | 5675 | 1558 |
| Sample 10 | 12 | 377 | 358 | 1089 | 4660 | 1379 | 33713 | 2087 | 3546 | 1287 |
| Sample 20 | 6 | 342 | 312 | 992 | 4138 | 1400 | 1801 | 1772 | 1458 | 805 |
| Sample 21 | 15 | 397 | 410 | 1160 | 4540 | 1666 | 3800 | 2370 | 5304 | 1515 |
| SRR15489009 | 12 | 367 | 388 | 1187 | 5194 | 1959 | 3194 | 1883 | 3186 | 594 |
| SRR15489011 | 11 | 364 | 380 | 1208 | 4655 | 1664 | 2860 | 1768 | 3055 | 449 |
| SRR15489017 | 14 | 377 | 416 | 1280 | 4991 | 2061 | 3380 | 1893 | 3150 | 665 |

We additionally tested Ram and Minimap2 execution times by mapping only one sequence to the whole database file. The execution time for both was around 1000 seconds, suggesting that the database parsing and indexing took that much time. Both tools could improve execution time by storing and loading preprocessed database indexes to the disk.

For Bracken, we analysed the running time and memory consumption for the database building procedure because it needed to be executed for every dataset independently since datasets had different average read lengths, a parameter required by Bracken. The abundance estimation script executed almost instantaneously.

We analysed the scalability of used tools on several different dataset sizes. The results are presented in Table 5. Even for the largest datasets, Ram was still, at most, around 10x slower than Kraken2, the fastest kmer-based tool. Although Centrifuge was the fastest tool when analysing execution times presented in Table 5, Kraken2 had the lowest execution times when tested on larger datasets. This happened because, for smaller datasets, index loading took up a large part of the execution time, and Centrifuge had the smallest database index. On larger datasets, where the actual sequence classification took a greater part of the execution time, Kraken2 outperformed Centrifuge.

All resource usage measurements were performed on a machine with sufficient disk space, 775 GB RAM and 256 virtual CPUs (2 x AMD EPYC 7662 64-Core Processor). Measurements were performed using 12 threads for 7 synthetic and 3 mock datasets and using 32 threads for 6 gut datasets. We cleared RAM Cache, File system buffer, and Swap space between runs.

## Discussion

In this manuscript, we analysed four categories of tools for microbial classifications: kmer-based tools, mapping-based tools, general-purpose long-read mappers and tools which use protein databases. In most of the tests, tools which use protein databases performed worse than other categories. However, it would be interesting to additionally test them using RNAseq data, especially considering that long reads technologies can sequence the whole transcripts.

Mapping-based tools apart from deSAMBA were much slower. They all required a lot of memory while rarely performing better than other tools. General purpose mappers, especially Minimap2 in the alignment mode, achieved the best results in most tasks. Ram achieved good results, and its low memory consumption makes it suitable for analysis on laptops. However, mappers were slower than kmer-based tools. Considering this, kmer-based tools (Kraken2 and Centrifuge) will still be the tools of choice in most scenarios, especially in differential studies.

An interesting aspect is the detection of present organisms. Most tools, especially those based on k-mers, overreported the number of present species. In their analysis, McIntyre et al. [37] found that different strategies, including abundance filtering, ensemble approach and tool intersection, help for short reads. Some tools, such as Kraken2, had an internal option for setting up the threshold for each read based on the number of find kmers. However, even these strategies could not completely eliminate false positives. A detailed assessment of various filtering and ensembling strategies is out of the scope of this paper, and it is a promising avenue for future research. Here, we show that setting up read thresholds reduces the number of false positives but can lead to a reduction in true positives. Moreover, longer reads help to increase accuracy. In parallel with our work, Portik et al [10] report the same. Finally, testing on mock communities showed that a lower error rate of PacBio reads might reduce false positives.

A special case in this analysis was PB_Zymo dataset. Differences between declared and calculated abundances for this tool are much larger than for other tools The same we noticed for all tools. Reasons for these might be in the upstream procedures, including sample preparation, DNA extraction and sequencing.

Although we didn’t know the ground truth for the real gut datasets, they helped us to understand the limitations and importance of carefully curated databases. For example, differences in calculated absolute abundances between tools were up to 50%, and all kmer-based tools didn’t report one species often found in the gut microbiome. This shows that using just one pipeline for sensitive experiments is probably insufficient. In addition, despite protein-based tools performing worse in this benchmark, it would be interesting to use them in the case when large amounts of reads are not associated with taxa in the nucleic database. We have found there are differences in taxa between nucleic and protein databases, so protein-based tools might indicate the presence of some taxa not contained in the nucleic database.

Analysis conducted on samples containing species not found in the database and real gut samples highlights the significant advantages of using meticulously curated databases. To ensure maximum fairness in this study, we created databases for each tested tool using the same input sequences. Created databases contain only complete sequences. For many taxa, only partial information was available. However, even among complete sequences, many unknown nucleotides or amino acids exist.

In this work, we used a measure we call the relative genome count abundance, which considers the length of reads and genome size. Using read counts and calculating ratios of genomic DNA for each species or counting genomes or cells is an open question. Recently, Sun et al [38] also warned of the importance of distinguishing these measures. It is important to note that the accuracy improved when we added read lengths to calculate the abundance for the ONT Zymo dataset. However, the results were slightly worse for mock communities sequenced by the PacBio HiFi technology. Investigating this in detail will be one of our next aims.

In the field of metagenomics, an ongoing challenge is how to effectively benchmark classification tools using metagenomic samples. In this study, we utilised real reads, mock communities, and varied ratios of present organisms in datasets to benchmark the performance of tools. Some researchers, such as McIntyre et al. [37], have used synthetic reads to create datasets that mimic the complexity of real samples. Although we also incorporated simulated datasets in our analysis, we acknowledge that these datasets may only partially replicate the complexity of real-world cases. However, most tools, including Minimap2, Ram, Kraken2, DeSamba, MEGAN-N, MEGAN_P, CLARK, CLARK-S, and Kaiju, analyse each read independently, and complex samples can be simulated by combining simple datasets linearly without significantly influencing the final results. Furthermore, we explored samples with logarithmic differences in species abundance. Our analysis revealed that different tools report varying species compositions for real gut datasets, and the true complexity of real samples remains incompletely understood. Therefore, we posit that a comprehensive study is needed to elucidate the differences between these two approaches to dataset preparation and their impact on the benchmark results.

This benchmark study shows that there is a discordance between tools, and we argue that there is room for improvement in all tools. A useful upgrade would be to provide some information about the confidence of whether the read belongs to a similar species or it doesn’t belong to any species in the database.

Since the existing deep learning models, such as DeepMicrobes [39] and BERTax [40], have been rarely used and primarily focuse on short reads, we did not include them in this benchmark. However, due to rapid progress in the usage of AI in all other fields and ample sequenced data, we believe that using deep learning methods, including those based on language models, will be one of the new avenues in this field.

## Conclusion

The assessment on real gut datasets shows that tools report a different number of species and different abundances. Therefore we deem that classifying microbial organisms is still an open problem.

Due to their speed, kmer-based tools are still the first option for most analyses. However, if the time is not too critical or there is not enough RAM, off-the-shelf mappers, including Minimap2 and Ram, are a good alternative which provides higher accuracy in most scenarios. Modern mappers use fewer kmers to calculate mapping candidate positions, which makes them faster. We believe that with further improvement in long-read sequencing technology, most methods will move to detect smaller numbers of kmers in combination with chaining matches. They will probably need an additional postprocessing step using methods such as the EM algorithm or careful filtering to reduce the number of false positives.

In this manuscript, we use relative genome count as an abundance measure, and we focused on datasets constructed using real reads from previous experiments, mock communities or real gut samples. However, we are aware of the used abundance measure and benchmark dataset preparation, and their alternatives have limitations. Hence, we recognise the need for further research in defining appropriate abundance measures and improving metagenomics benchmark dataset construction techniques in the field.

Although there are just a few deep-learning methods for microbial classification, we expect a sharp increase in their number in the following months and years.

The results obtained from the analysis of real gut samples clearly indicate that using different strategies and database types may be necessary to accurately determine microbial content. Furthermore, our findings strongly support the notion that there is ample room for improvement in systematically incorporating key microbial species into existing databases containing reference genomes and protein sequences. We believe that advancements in long-read sequencing technologies will greatly facilitate and enhance this process.

## Methods

This chapter describes the tools and assessment measures used and how we constructed the test datasets and databases.

## Databases

Each classification tool comes with a prebuilt default database and instructions on building and using a custom database. We created a database for each tool based on the same set of organisms to remove bias related to default databases. We used NCBI-NT and NCBI-NR databases for creating protein and nucleotide databases used in this work We downloaded bacteria, archaea, and human sequences whose *assembly_level* property in the *assembly_summary.txt* file (downloaded from the RefSeq directory) is “Complete Genome” or “Chromosome”. In the following text, we refer to the two databases that have been created as the protein and nucleotide databases.

The list of all the downloaded files and all the sequence ids of the databases can be found in the GitHub link: (https://github.com/lbcb-sci/MetagenomicsBenchmark).

Genome sequences were downloaded on April 5^th^ 2020, along with the taxonomy files nodes.dmp and names.dmp. This approach allowed the tools to be tested independently of the content of their default database. The details of how each database index was created for every tool are presented in Supplementary Materials 3.

The main part of the tests was performed on a database constructed for each tool from the same set of sequences: protein databases created from NCBI-NR database for tools using a protein database and nucleotide database created from NCBI-NT database for tools using a nucleotide database. We used all bacterial and archaeal genomes, plus the human genome. The nucleotide database contains 9044 tax ids, while the protein contains 11435 tax ids. There are 552 tax ids in the protein database that are outside the nucleotide database, and there are 2943 tax ids that are in the protein database but not in the nucleotide database. It is important to highlight that only one species (*Helcococcus kunzii* – 40091) is present in mock datasets and the protein database but not in the nucleotide database. Supplementary Table 10 shows a list of dataset species not present in databases. The taxonomy data was downloaded from this website: https://ftp.ncbi.nlm.nih.gov/pub/taxonomy/accession2taxid/ and genome sequences were Downloaded from this website: https://ftp.ncbi.nlm.nih.gov/genomes/refseq/.

## Test datasets

To have realistic sequencing datasets while retaining control of our mock communities’ exact content and building the ground truth, we constructed in silico datasets by mixing real reads from isolated, sequenced species. Data was downloaded from multiple sources (details in Supplementary Table 7), including the European Nucleotide Archive (ENA [41]) and the National Center for Biotechnology Information (NCBI [42]). This in-silico approach provides ground truth and great flexibility to create diverse datasets while offering real reads with their natural errors and length variance. Most datasets contained around 100,000 reads, allowing all tools to classify them within a week. We varied the proportion of species, some with even distributions, some with decreasing ratios with as little as five reads for one species (PB4 dataset). Seven test datasets were synthesised with the following composition: two ONT, four PacBio and one negative dataset containing PacBio and randomised reads.

– **ONT1**: 18 bacterial species with a percentage of reads varying from 18% down to 0.01%.
– **ONT2**: Human (about 4000 reads) + 7 bacteria, 10,000 reads each.
– **PB1**: 10 bacteria, 10% each (including two strains of *Escherichia coli*).
– **PB2**: 20% human reads, 20% fruit fly (*Drosophila melanogaster*), 10% archaea (*Methanocorpusculum labreanum* Z), and ten bacteria, varying from 10% to ∼1%.
– **PB3**: 99% human reads, plus two bacteria: 0.9% *E. coli* and 0.1% *Staphylococcus aureus*.
– **PB4**: 46 bacterial species with the percentage of reads varying from 10% to 0.005%.
– **PB1+NEG**: It contains all the reads from PB1 datasets with additional 20000 “randomised” reads that should not be assigned to any organism. Randomised reads were obtained by shuffling the human genome (GRCH38.p7) using *esl-shuffle* script from the hmmer3 [43] package (version 3.3.2) as described by Lindgreen et al. [44].

All datasets that do not contain human reads were mapped to the human reference with minimap2 to check if there are human reads’ contaminations. No sequences that belong to non-human species mapped to the human genome with a significant quality.

In addition to synthetic datasets, the tools were tested on three real datasets obtained by sequencing mock metagenomic communities. The results reported by the tested tools were used to calculate abundances and compared to standard specifications obtained from manufacturer pages.

– **ONT_Zymo**: obtained by GridION sequencing of a Zymo Community Standard, consists of eight bacteria and two yeasts with the expected abundance varying from 0.37% to 21.6% (downloaded from Loman Labs https://lomanlab.github.io/mockcommunity/).
– **PB_ATCC**: obtained by PacBio HiFi sequencing of an ATCC MSA-1003 standard (20 Strain Staggered Mix Genomic Material), consists of 20 different bacterial species with the expected abundance varying from 0.02% to 18% (download from NCBI archive, SRA run identifier: SRR11606871).
– **PB_Zymo**: obtained by PacBio HiFi sequencing of a Zymo D6331 Gut Microbiome Standard, consisting of 16 bacteria and one yeast, with the expected abundance varying from 0.0001% to about 20% (download from NCBI archive, SRA run identifier: SRR13128014). However, for this dataset, the results obtained by all tools differed significantly from the specification.

Finally, the tools were tested on six real human gut microbiome datasets. Three datasets were produced as a part of the CaPES study at the Genome Institute of Singapore and published on ENA under the project ID PRJEB29152 (Bertrand et al. 2019). Datasets used for our test:

– Sample10 – 11.9 GB
– Sample20 – 6.4 GB
– Sample21 – 14.8 GB

All three samples were sequenced using an R9.5 spot-on flowcell. Sample 10 was basecalled using Albacore, while samples 20 and 21 were basecalled using Guppy v0.3.0 for live 1D basecalling.

The other three datasets were generated as a part of a clinical trial by Siolta Therapeutics [6]. All generated data is available in the NCBI database under the BioProject accession number PRJNA754443. All long-read datasets were sequenced using a PacBio Sequel II. Datasets used in our testing:

– SRR15489009 – 12GB
– SRR15489011 – 11GB
– SRR15489017 – 14GB

## Tools

Tested tools can be classified into:

– k-mer based: Kraken2 (v2.0.8), Centrifuge (v1.0.4), CLARK (v1.2.6.1) and CLARK-S
– mapping-based: MetaMaps (v0.1), MEGAN-LR (v6.18.5) (using a nucleotide database, named MEGAN-N), deSAMBA (v1.1.12),
– General purpose long-read mappers: Minimap2 (v2.18) and Ram (v2.1.1)
– tools which use protein databases: Kaiju (v1.8.2) and MEGAN-LR (v6.18.5) (named MEGAN-P)

Since Kraken2 usually uses Bracken for the calculation of abundances, we included it in the analysis.

Tools start with the initial assignment of reads to genomes using in-advance-prepared databases of known organisms. Once when all reads are assigned, various methods are used to fine-tune the classification using information from assigned reads and taxonomy trees. The most popular postprocessing approaches are Expectation-Maximization (EM) estimation (MetaMaps, Centrifuge), Bayesian estimation (Bracken) and read assignment using the least common ancestor approach (MEGAN-LR, Kraken2).

The initial assignment of reads is based on aligning reads to a database of determined genomes. Aligning (Figure 5) might be divided into three steps: (1) Searching for exact or approximate matches of short substrings of length k (kmers) or longer in a previously prepared index which contains a list of kmers from genomes, (2) Chaining kmer matches into a sequence, scoring the sequence, finding approximate positions of a read in a genome (mapping), and choosing the best genome candidates (3) Alignment of a read and candidate genomes using exact dynamic programming algorithm. While kmer-based tools use only the first step, mapping-based tools use the first and second or all three. Each additional step adds to accuracy but significantly increases the running time.

**Figure 5.**
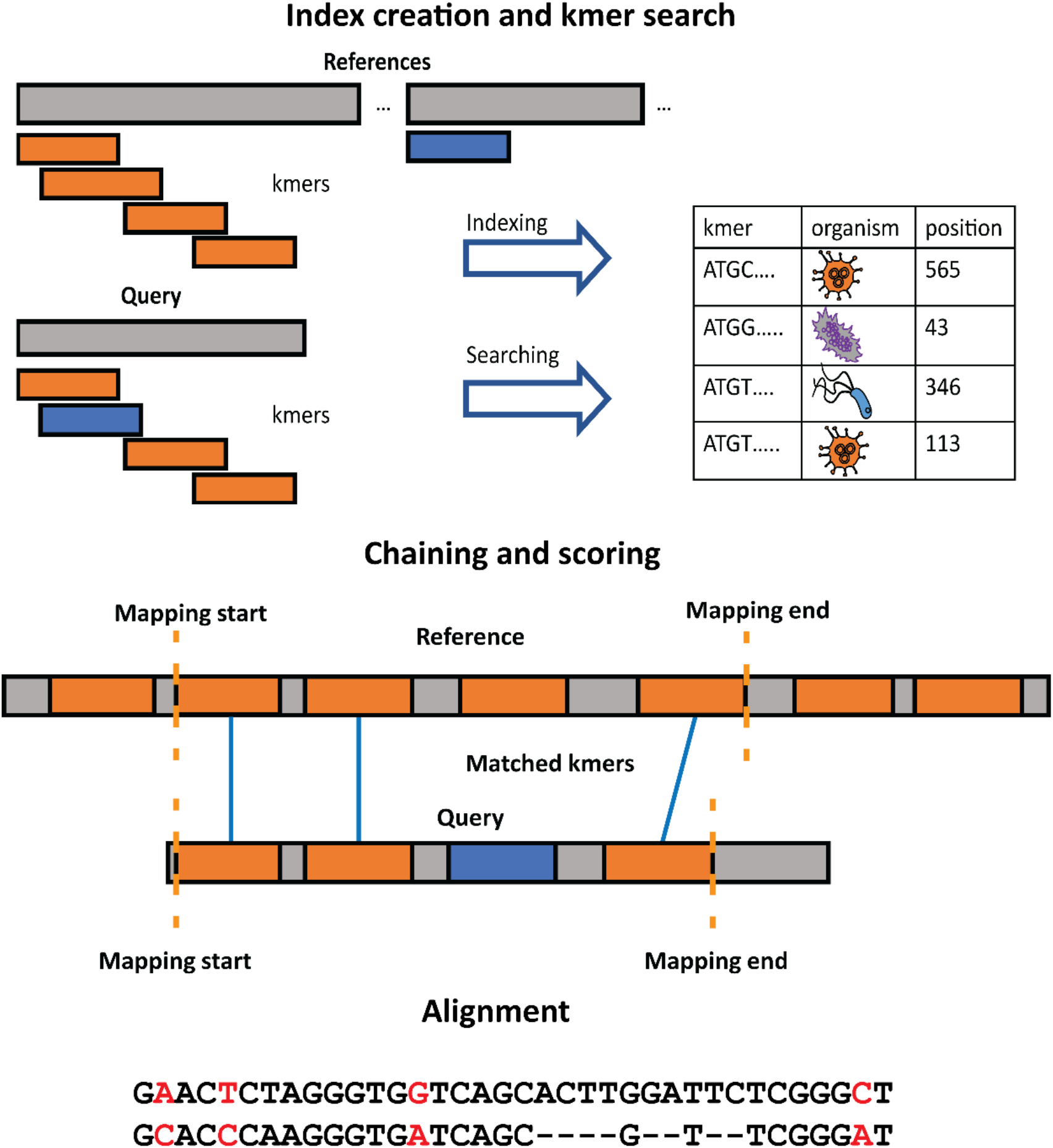
Read alignment. Read alignment consists of three steps (1) Indexing and kmer search, (2) Chaining and scoring (3) Alignment. Kmer-based tools use only the first step, and usually, they do not care about the position in the genome. Mapping-based tools use the first and second steps, which increase accuracy but last much longer. The alignment step provides the exact alignment and the score but increases the running time.

Usually, kmers are of a fixed size. The original approach used all sliding windows of size k in a sequence. This might lead to high accuracy, but it is too slow. Therefore, modern tools usually use just a few discriminative kmers per genome or choose a lexicographically smallest kmer in a window of w consecutive kmers, i.e. minimiser [45].

The outputs of various tools were processed to obtain read-level classifications and an abundance of various species in a sample. Short descriptions and versions of each tool are available in Supplementary Materials 1. Specific parameters and scripts used to run each tool are given in Supplementary Materials 2.

Furthermore, we could not successfully run our version of the MEGAN–N pipeline on the PB3 synthetic dataset and on PB_Zymo and PB_ATCC real datasets. In the case of the PB3 dataset, the mapping phase using the LAST aligner would go on for a week, and after that, the CPU and memory usage would drop down to almost zero, but the process would not be complete. Output produced in that way was corrupted and could not be used for testing. After three trials, we decided to drop the results. For PB_Zymo and PB_ATCC datasets, the LAST aligner produced a huge *MAF* file with correct alignments, which we could not convert to an alignment out file (.*DAA*). This resulted in no classified reads. Huge MAF files and long running times were also a reason why we did not run MEGAN-N on larger datasets. MEGAN-P uses a protein database, produces significantly smaller *MAF* files, runs faster, uses less RAM and is able to complete most tests. MEGAN-P did not finish on the real gut microbiome datasets SRR15489009 and SRR15489017. Although *MAF* files produced by the LAST aligner seemed correct, the script used to convert *MAF* files into *DAA* files needed for further analysis produced a file with the data for only the first read.

Minimap2 and Ram are not intended for metagenomics classifications and often print several mapping results for a single sequence. Therefore we classify each sequence from the *PAF* output files using a harmonic mean:

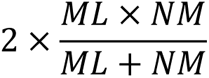

where the *ML (mapping length)* and *NM* (*number of matches)* are found in each row of the *PAF* file. For the *SAM* output files with alignment, the best classification for each sequence was determined by the highest alignment score. In the case of ties, the first alignment was taken as correct. This work aims not to maximise mappers’ performance but to demonstrate their suitability for metagenomic classification. Detailed analysis of ties presented in Supplementary Table 8 shows that more careful handling of ties might lead to further improvement in read classifications.

## Genome lengths

We acquired average genome lengths for species used in abundance calculation from the NCBI website: https://www.ncbi.nlm.nih.gov/genome/?term=<species_name>. We scraped the NCBI website for the “median total length (Mb)” string to get the genome length of the species. For species that were unsuccessfully scraped, we acquired the genome length from the same website manually.

## Testing procedures

The tools’ output was processed to obtain percentages of DNA reads and species’ abundances in the sample. We evaluated the correctness of DNA read classification at species and genus level, i.e., only classifications that were assigned to a tax id which belongs to the species or lower level were used in the species-level analysis; and only classifications assigned to the genus or a lower level were used in the genus-level analysis. Outputs of the tools, which contained classification of reads to taxons, were processed. Taxonomic ids and ranks were extracted from the nodes.dmp file downloaded from the NCBI website.

## Read-level classification

To evaluate the quality of read-level classification, we first calculated four basic values:

– True positives (TP): the number of reads that were classified to a correct species.
– False positives (FP): the number of reads that were classified as an incorrect species.
– True negatives (TN): the number of reads that remained unclassified and belonged to an organism not present in the database.
– False negatives (FN): the number of reads that remained unclassified but belonged to an organism present in the database.

These four values were then used to calculate more complex and valuable evaluation metrics. The first metric used is classification accuracy – the percentage of reads that were correctly classified.

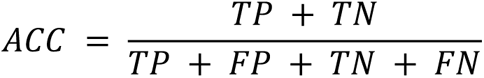

Global accuracy score was calculated for each read regardless of which organism it belonged to, only based on whether it was classified as TP, TN, FP or FN. As such, it did not reflect the proportion of each species in the sample. Incorrectly classified reads belonging to very abundant organisms will have the same effect as those belonging to very low abundance organisms. Because it is less biased towards larger classes, we used the F1 macro average score: we calculated the F1 score for each class (organism in the sample) separately and averaged them. We prefer this measure to the global F1 score, calculated across all classes. F1 macro average gives an equal impact on the score for each organism in the sample independently of its abundance.

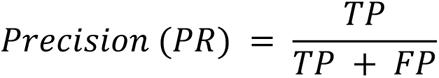

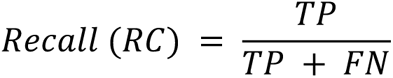

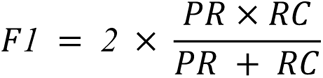

Because the F1 score is zero for classes not in the database (as the number of true positives is zero), those classes were omitted from the calculation.

### Abundance

Abundance represents the percentage of genomes of a specific taxon in the sample. Abundances calculated by benchmarked tools significantly differ due to differences in definitions and calculations. Here we calculated abundances for all tools using so-called relative genome count abundance measure. In addition to genome sizes in the calculation, we added read lengths due to the skewed and wide distribution of ONT reads. We calculated the abundance of a species as the sum of the lengths of assigned reads divided by its average genome length in the database. Obtained values were normalised in a manner that the total sum of abundances is 1. We calculated C_i_, a sequenced coverage of genome *i*, and A_i_, a relative abundance of genome *i* as

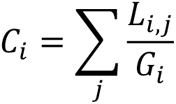

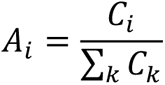

where *L_i,j_* is the length of a read *j* assigned to genome *i, G_i_* length of genome *i*, and *C_k_* is the sequenced coverage that belongs to any of the reported genomes.

We use L1 norm for measuring the accuracy of predicted abundances of taxa in a sample at rank *r.* The L1 norm is given by

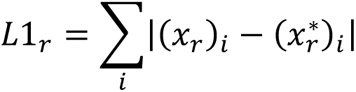

where *x*_%_ and *x_r_*\* are true and predicted abundances, respectively. We choose L1 instead of L2 because the former is less sensitive to outliers.

Genome lengths were obtained from the NCBI web page. Abundance estimations were assessed on species or lower-level classifications, considering all higher-level taxa classifications incorrect.

## Organisms detection

Although some benchmarks, including [37], used precision and recall as measures for assessing organism detection, we decided to show only the number of true positives and the total number of reported organisms. This helped us to make a similar assessment for all datasets, including those for real gut datasets where we reported only the number of detected organisms.

## Resource usage

Processor time and RAM usage were measured using a server with 775 GB RAM and 256 virtual CPUs (2 x AMD EPYC 7662 64-Core Processor). Tools were run using 12 threads for 7 synthetic and 3 mock datasets and 32 threads for 6 gut datasets. The measurements were taken using a fork of cgmemtime (https://github.com/isovic/cgmemtime) modified to write its output to a file.

## Declarations

### Ethics approval and consent to participate

Not applicable.

## Consent for publication

Not applicable.

## Availability of data and materials

Python scripts used to generate all figures and tables in this study, and all supplementary texts and figures are available via the Github repository https://github.com/lbcb-sci/MetagenomicsBenchmark. Datasets, supporting files, analysis results and reports are available via Zenodo repository https://doi.org/10.5281/zenodo.7213115.

## Authors’ contributions

M.Š. and N.N. conceived the project. K.K. and S.R. prepared the datasets. J. M. and K. K. ran the tests and developed auxiliary scripts. K.K. and J.M. planned and conducted the data analysis. M.Š, K.K and J. M. wrote the manuscript. All authors read and approved the final manuscript.

## Acknowledgements

NA

## References

1. Quince C, Walker AW, Simpson JT, Loman NJ, Segata N. Shotgun metagenomics, from sampling to analysis. Nat Biotechnol. 2017;35:833–44.

2. McFall-Ngai M, Hadfield MG, Bosch TCG, Carey HV, Domazet-Lošo T, Douglas AE, et al. Animals in a bacterial world, a new imperative for the life sciences. Proc Natl Acad Sci U S A. 2013;110:3229–36.

3. Hamady M, Knight R. Microbial community profiling for human microbiome projects: Tools, techniques, and challenges. Genome Res. 2009;19:1141–52.

4. Bertrand D, Shaw J, Kalathiyappan M, Ng AHQ, Kumar MS, Li C, et al. Hybrid metagenomic assembly enables high-resolution analysis of resistance determinants and mobile elements in human microbiomes. Nat Biotechnol. Springer Science and Business Media LLC; 2019;37:937–44.

5. Chen L, Zhao N, Cao J, Liu X, Xu J, Ma Y, et al. Short– and long-read metagenomics expand individualized structural variations in gut microbiomes. Nat Commun. 2022;13:3175.

6. Gehrig JL, Portik DM, Driscoll MD, Jackson E, Chakraborty S, Gratalo D, et al. Finding the right fit: evaluation of short-read and long-read sequencing approaches to maximize the utility of clinical microbiome data. Microb Genom [Internet]. 2022;8. Available from: http://dx.doi.org/10.1099/mgen.0.000794

7. Pearman WS, Freed NE, Silander OK. Testing the advantages and disadvantages of short– and long-read eukaryotic metagenomics using simulated reads. BMC Bioinformatics. 2020;21:220.

8. Nicholls SM, Quick JC, Tang S, Loman NJ. Ultra-deep, long-read nanopore sequencing of mock microbial community standards. Gigascience [Internet]. 2019;8. Available from: http://dx.doi.org/10.1093/gigascience/giz043

9. Leidenfrost RM, Pöther D-C, Jäckel U, Wünschiers R. Benchmarking the MinION: Evaluating long reads for microbial profiling. Sci Rep. 2020;10:5125.

10. Portik DM, Brown CT, Pierce-Ward NT. Evaluation of taxonomic classification and profiling methods for long-read shotgun metagenomic sequencing datasets. BMC Bioinformatics. Springer Science and Business Media LLC; 2022;23:541.

11. Wood DE, Lu J, Langmead B. Improved metagenomic analysis with Kraken 2. Genome Biol. 2019;20:257.

12. Lu J, Breitwieser FP, Thielen P, Salzberg SL. Bracken: estimating species abundance in metagenomics data. PeerJ Comput Sci. PeerJ; 2017;3:e104.

13. Kim D, Song L, Breitwieser FP, Salzberg SL. Centrifuge: rapid and sensitive classification of metagenomic sequences. Genome Res. 2016;26:1721–9.

14. Ounit R, Wanamaker S, Close TJ, Lonardi S. CLARK: fast and accurate classification of metagenomic and genomic sequences using discriminative k-mers. BMC Genomics. 2015;16:236.

15. Ounit R, Lonardi S. Higher classification sensitivity of short metagenomic reads with CLARK-S. Bioinformatics. 2016;32:3823–5.

16. Dilthey AT, Jain C, Koren S, Phillippy AM. Strain-level metagenomic assignment and compositional estimation for long reads with MetaMaps. Nat Commun. 2019;10:3066.

17. Huson DH, Albrecht B, Bağcı C, Bessarab I, Górska A, Jolic D, et al. MEGAN-LR: new algorithms allow accurate binning and easy interactive exploration of metagenomic long reads and contigs. Biol Direct. 2018;13:6.

18. Li G, Liu Y, Li D, Liu B, Li J, Hu Y, et al. Fast and Accurate Classification of Meta-Genomics Long Reads With deSAMBA. Front Cell Dev Biol. 2021;9:643645.

19. Li H. Minimap2: pairwise alignment for nucleotide sequences. Bioinformatics. 2018;34:3094–100.

20. Vaser R, Šikić M. Time– and memory-efficient genome assembly with Raven. Nature Computational Science. Nature Publishing Group; 2021;1:332–6.

21. Menzel P, Ng KL, Krogh A, Riesenfeld C, Schloss P, Handelsman J, et al. Fast and sensitive taxonomic classification for metagenomics with Kaiju. Nat Commun. Nature Publishing Group; 2016;7:11257.

22. Ainsworth D, Sternberg MJE, Raczy C, Butcher SA. k-SLAM: accurate and ultra-fast taxonomic classification and gene identification for large metagenomic data sets. Nucleic Acids Res. 2017;45:1649–56.

23. Truong DT, Franzosa EA, Tickle TL, Scholz M, Weingart G, Pasolli E, et al. MetaPhlAn2 for enhanced metagenomic taxonomic profiling. Nat Methods. Nature Publishing Group; 2015;12:902–3.

24. Luo C, Knight R, Siljander H, Knip M, Xavier RJ, Gevers D. ConStrains identifies microbial strains in metagenomic datasets. Nat Biotechnol. 2015;33:1045–52.

25. Hong C, Manimaran S, Shen Y, Perez-Rogers JF, Byrd AL, Castro-Nallar E, et al. PathoScope 2.0: a complete computational framework for strain identification in environmental or clinical sequencing samples. Microbiome. BioMed Central; 2014;2:33.

26. Breitwieser FP, Baker DN, Salzberg SL. KrakenUniq: confident and fast metagenomics classification using unique k-mer counts. Genome Biol. 2018;19:198.

27. Ahn T-H, Chai J, Pan C. Sigma: strain-level inference of genomes from metagenomic analysis for biosurveillance. Bioinformatics. 2015;31:170–7.

28. Marcelino VR, Clausen PTLC, Buchmann JP, Wille M, Iredell JR, Meyer W, et al. CCMetagen: comprehensive and accurate identification of eukaryotes and prokaryotes in metagenomic data. Genome Biol. 2020;21:103.

29. Martin DMA, Berriman M, Barton GJ. GOtcha: a new method for prediction of protein function assessed by the annotation of seven genomes. BMC Bioinformatics. 2004;5:178.

30. Buchfink B, Xie C, Huson DH. Fast and sensitive protein alignment using DIAMOND. Nat Methods. 2014;12:59–60.

31. Fan J, Huang S, Chorlton SD. BugSeq: a highly accurate cloud platform for long-read metagenomic analyses. BMC Bioinformatics. 2021;22:160.

32. Vacca M, Celano G, Calabrese FM, Portincasa P, Gobbetti M, De Angelis M. The Controversial Role of Human Gut Lachnospiraceae. Microorganisms [Internet]. 2020;8. Available from: http://dx.doi.org/10.3390/microorganisms8040573

33. Karcher N, Pasolli E, Asnicar F, Huang KD, Tett A, Manara S, et al. Analysis of 1321 Eubacterium rectale genomes from metagenomes uncovers complex phylogeographic population structure and subspecies functional adaptations. Genome Biol. 2020;21:138.

34. Alpizar-Rodriguez D, Lesker TR, Gronow A, Gilbert B, Raemy E, Lamacchia C, et al. Prevotella copri in individuals at risk for rheumatoid arthritis. Ann Rheum Dis. 2019;78:590– 3.

35. Wang L, Christophersen CT, Sorich MJ, Gerber JP, Angley MT, Conlon MA. Increased abundance of Sutterella spp. and Ruminococcus torques in feces of children with autism spectrum disorder. Mol Autism. 2013;4:42.

36. Ferreira-Halder CV, Faria AV de S, Andrade SS. Action and function of Faecalibacterium prausnitzii in health and disease. Best Pract Res Clin Gastroenterol. 2017;31:643–8.

37. McIntyre ABR, Ounit R, Afshinnekoo E, Prill RJ, Hénaff E, Alexander N, et al. Comprehensive benchmarking and ensemble approaches for metagenomic classifiers. Genome Biol. 2017;18:182.

38. Sun Z, Huang S, Zhang M, Zhu Q, Haiminen N, Carrieri AP, et al. Challenges in benchmarking metagenomic profilers. Nat Methods. Springer Science and Business Media LLC; 2021;18:618–26.

39. Liang Q, Bible PW, Liu Y, Zou B, Wei L. DeepMicrobes: taxonomic classification for metagenomics with deep learning. NAR Genom Bioinform. 2020;2:lqaa009.

40. Mock F, Kretschmer F, Kriese A, Böcker S, Marz M. BERTax: taxonomic classification of DNA sequences with Deep Neural Networks [Internet]. bioRxiv. 2021 [cited 2022 Oct 8]. p. 2021.07.09.451778. Available from: https://www.biorxiv.org/content/biorxiv/early/2021/07/10/2021.07.09.451778

41. Leinonen R, Akhtar R, Birney E, Bower L, Cerdeno-Tárraga A, Cheng Y, et al. The European Nucleotide Archive. Nucleic Acids Res. 2011;39:D28–31.

42. Sayers EW, Barrett T, Benson DA, Bolton E, Bryant SH, Canese K, et al. Database resources of the National Center for Biotechnology Information. Nucleic Acids Res. 2011;39:D38–51.

43. Eddy SR. Accelerated profile HMM searches. PLoS Comput Biol. Public Library of Science (PLoS); 2011;7:e1002195.

44. Lindgreen S, Adair KL, Gardner PP. An evaluation of the accuracy and speed of metagenome analysis tools. Sci Rep. Springer Science and Business Media LLC; 2016;6:19233.

45. Roberts M, Hayes W, Hunt BR, Mount SM, Yorke JA. Reducing storage requirements for biological sequence comparison. Bioinformatics. 2004;20:3363–9.

