## Supplementary materials for "Comprehensive benchmarking of metagenomic classification tools for long-read sequencing data"

^#^To whom correspondence should be addressed.

Supplementary materials

### Supplementary Materials - Contents

**Supplementary Materials 1 - Tools**: Description of tested tools.

**Supplementary Materials 2 - Tools usage**: Detailed explanation of how each tool was run, with used command-line instructions.

**Supplementary Materials 3 - Database creation**: Detailed explanation of how a database was built for each tool, based on sequences from NCBI-NT or NCBI-NT database, with command-line instructions.

**Supplementary Figure 1**: Read level classification F1 macro average, comparison between species and genus level classification.

**Supplementary Figure 2**: In-sample abundance error boxplots for all datasets.

**Supplementary Figure 3:** In-sample abundance error boxplots for mock community datasets - comparison between read counts and sum of reads’ lengths.

**Supplementary Table 1**: Total absolute abundance estimation error using 30% of longest reads and using all reads.

**Supplementary Table 2**: Resource usage.

**Supplementary Table 3**: Percentage of reads classified for all datasets and tools.

**Supplementary Table 4**: Number of species detected for all real datasets and all tools.

**Supplementary Table 5**: Composition of Zymo and ATCC standards.

**Supplementary Table 6**: Genome lengths of species having more than 1% of reads classified for three real datasets.

**Supplementary Table 7**: Isolate datasets used for testing dataset construction.

**Supplementary Table 8**: Tie handling while processing mappers output.

**Supplementary Table 9**: Abundance estimation error for low abundant species.

**Supplementary Table 10**: Analysis of the difference between relative cell count and read count calculated abundances for dataset PB_ATCC

**Supplementary Table 11:** Abundance results for SRR15489009(PacBio) dataset.

### Supplementary Material 1 - Tools

**Kraken2**

Kraken is a tool that assigns taxonomic labels to DNA sequences using exact alignments of k-mers^10^. In this study, we used Kraken2, an improved version of Kraken, which reduces memory usage. The Kraken2 report file contains summarised hierarchical percentages of taxa abundances in the dataset, while the classification file contains the unique classification of each read to an assigned taxa id. We used the latter to calculate the percentages of reads assigned to the corresponding taxa id. Kraken (v 2.0.8) was run with default settings.

**Bracken**

Bracken estimates species abundances in metagenomics samples by probabilistically re-distributing reads in the taxonomic tree by reassigning sequences assigned to the nodes above the species level down to the species nodes.

**Centrifuge**

Centrifuge is a metagenomic classification tool that uses an indexing scheme based on the Burrows-Wheeler transform (BWT) and the FM index. Centrifuge (v 1.0.4) was run with default settings, but with the parameter k=1 to set the tool to classify each read to only one taxon.

**CLARK**

CLARK builds an extensive index of discriminative k-mers for all target sequences, and an object (sequencing read) is assigned to a target sequence with which it shares the highest number of k-mers. CLARK (v 1.2.6.1) was run with default settings.

**CLARK-S**

CLARK-S builds on top of CLARK and increases its classification sensitivity by allowing mismatches between read's and reference's shared k-mers in a limited number of positions while maintaining the requirement for k-mers to be discriminative.

**MetaMaps**

MetaMaps (v 0.1) performs classification in two steps. In the first step, possible mapping locations are generated using an approximate mapping algorithm based on minimizers. In the second step, mapping locations are scored probabilistically, and a total sample composition is estimated using an EM algorithm.

**MEGAN-LR**

MEGAN is a comprehensive toolbox for interactively analyzing microbiome data. It uses an LCA-based algorithm for taxonomic binning and an interval-tree-based algorithm for functional binning. In 2018, it was upgraded to work with long reads - MEGAN-LR.

The MEGAN-LR pipeline uses the LAST aligner to align long reads to a protein database, and an auxiliary script to convert alignments to the .DAA format and the MEGAN application to analyze the data.

MEGAN 6 community edition version 6_18_5 with LAST aligner version 2.27.1 was used. Although the MEGAN-LR pipeline was originally used with the NR protein database, to be able to have the same database for all tools, we also used it on the NT nucleotide database. Reads were first mapped to the reference database (protein or nucleotide) using the LAST aligner. The .MAF file produced was converted to .DAA format using the maf2daa tool. A .DAA file was prepared for import to MEGAN-LR, using the daa-meganizer tool with the option -lg for long reads. Finally, read counts per species were extracted in CSV format using MEGAN command line mode.

Constructing a custom database for MEGAN-LR is actually creating the LAST index on the corresponding set of reference genomes. The instructions used were obtained from the LAST tutorial page.

**Minimap 2**

Minimap2 is the state-of-the-art mapping tool for long reads. The details about how Minimap2 (v 2.18) was used can be found in the Supplementary Materials 2.

**Ram**

Ram is a rewritten Minimap tool with the difference that it uses a reduced number of minimizers. Only lexicographically smallest |read|/kmer_size minimizers are picked in both the index and the query. The details about how Ram (v 2.1.1) was used can be found in Supplementary Materials 2.

Since the Ram outputs the results in the same format as Minimap2, PAF format, the classification results are obtained similarly as for Minimap2.

**Kaiju**

Kaiju (v 1.8.2) is a metagenome classifier which employs a search strategy which finds maximal exact matching substrings between query and database on the protein level, using the
Burrows-Wheeler transform.

**deSAMBA**

deSAMBA (v 1.1.12) is a long read classification tool that uses Unitig-BWT data structure to index the unitigs of the de Bruijn graph of the reference sequences and finds similar blocks between reads and references through the index.

### Supplementary Material 2 - Tools usage

For every tool, classification of the sequences to the tax ids were obtained. Only the sequences classified to the species or lower taxonomy level were used in the species classification analysis, and only sequences classified to the genus or lower lever were used in the genus classification analysis.

2.1. Kraken2

Kraken2 was run with the default settings.

> kraken2 --report <*report_file>* --db <*database_path>* <*dataset_path>* > <*classification_file>*

The resulting Kraken's database index takes around 42 GB of disk space.

Kraken's *<classification_file>* file, which is in standard kraken output format, was parsed to obtain the sequence classifications to the species by extracting sequence ids and tax ids from each row.

2.2. Bracken

Bracken was run with the default settings. Bracken's index was built for every dataset because every dataset has a different average read length. The command used to build Bracken's index:

> bracken-build -d <*kraken_database>* -l <*average_read_length>* -x <*kraken_installation_path>*

Bracken's estimation of read counts, denoted as abundance in Bracken paper, was done with following command:

> est_abundance.py -i <*kraken_report_file>* -k <*bracken_database_file>* -o <*bracken_results>*

2.3. Centrifuge

Centrifuge was run with default settings, but with the parameter k=1 to set the tool to classify each read to only one taxon. The command used was:

> centrifuge -x <*path_to_database> -k 1* -f <*path_to_dataset>* -S <*classification_file>* --report-file <*report_file>*

Centrifuge's index takes around 35 GB of space.

Centrifuge's <classification_file> file, which is in standard centrifuge output format, also contained sequence ids and tax ids in each row of the file, which were extracted to obtain the sequence classifications to the species.

#### 2.4. CLARK and CLARK-S

CLARK was run with the default settings. The databases for classification are chosen with the following command:

> set_targets.sh *<path_to_database>* custom

The classification is done with the following command:

> classify_metagenome.sh -O <*path_to_dataset>* -R <*classification_file>*

If it has not been previously created, this command will build the database index which takes around 125 GB of disk space.

CLARK-S additional index files, that take 228 GB of disk space, were generated with the command:

> buildSpacedDB.sh

Clark's <classification_file> file, which is in the format *Object_ID,Length,Assignment*, contains sequence classifications to the tax ids in each line that were extracted to obtain sequence classifications.

#### 2.5. MetaMaps

MetaMaps was used to classify reads with the following commands:

> metamaps mapDirectly --all -r <*path_to_database>* -q <*path_to_dataset>* -o <*classification_file_prefix>*

> metamaps classify --mappings <*classification_file_prefix>* --DB <*path_to_database>*

First command produces mappings of reads to reference genomes in the database. The second command then calculates the final classification of reads to tax id's.

MetaMaps's index takes around 66 GB of disk space.

MetaMaps produces several output files as the result of the classification. The file <classification_file_prefix>.EM.reads2Taxon.krona that contains final mappings of sequence ids to the tax ids in each row was used to obtain sequence classifications to species.

#### 2.6. MEGAN-LR

MEGAN6 is a comprehensive toolbox for interactively analyzing microbiome data ^16^. It uses an LCA-based algorithm for taxonomic binning and an interval-tree based algorithm for functional binning. In 2018, it was upgraded to work with long reads - Megan-LR ^17^.

The MEGAN-LR pipeline uses the LAST aligner to align long reads to a protein database, a script to convert alignments to the .DAA format and the MEGAN application to analyze the data.

MEGAN6 community edition version 6_18_5 with Last aligner version 2.27.1 was used.

MEGAN was used with both databases, the NT nucleotide database and the NR protein database. Reads were first mapped to the reference database using the LAST aligner. The .MAF file produced was converted to .DAA format using the maf2daa tool. A .DAA file was prepared for import to MEGAN, using the daa-meganizer tool with the option *-lg* for long reads. Finally, read counts per species were extracted in CSV format using MEGAN command line mode with the following script:

> open file='/mnt/data/megan-test/test_dataset_NR.daa';

> update;

> select nodes=all;

> export what=CSV format=readName_to_taxonId separator=tab counts=assigned file='/home/kkrizanovic/metabenchmark/results/Mock_100000-bacteria-l1000-q10_2BacArHum-RN2TAX.csv';

> quit;

Regarding the custom database creation, the MEGAN-LR pipeline uses the LAST aligner for mapping, and constructing a custom database is actually creating a LAST index on the corresponding set of reference genomes. The instructions used were obtained from the LAST tutorial page: <http://last.cbrc.jp/doc/last-tutorial.html>.

#### 2.7. Minimap2

Minimap2 was used to classify reads with the following command:

> minimap2 -ax <mode> --secondary=no --sam-hit-only <*path_to_database>* <*path_to_dataset>* > <*classification_file>*

The tool was run with alignment generation (-ax) and without alignment generation (-x). Depending on the dataset type the tool was run in different modes: *map-pb*, *map-ont* and *map-hifi*. Finally, the output files, *sam* file when using the alignment and *paf* file when not using the alignment, were parsed to obtain the classifications of sequences to the species.

#### 2.8. Ram

Ram was ran with the following command:

> ram --minhash <*path_to_database>* <*path_to_dataset>* > <*classification_file>*

2.9 Kaiju

Kaiju was run with the following command:

> kaiju -t <path_to_nodes_file> -f <path_to_database_index_file> -i <path_to_dataset> -o <path_to_output_file>

<path_to_nodes_file> is the nodes.dmp file downloaded with the protein database.

Kaiju's index takes around 88GB.

The output file contains classifications for each read where every line contains read_id to tax_id mapping and the C/U letter at the start of the line indicating if that read was classified.

2.10 deSAMBA

deSAMBA was run with the following command:

> deSAMBA classify -r 0 <path_to_database_index> <path_to_dataset> -o <path_to_output_file>

The tool was run with the -r parameter to not print any secondary alignments so that every read has only the best classification.

deSAMBA index takes around 146GB.

Output file was in the *sam* format and the best classification for each read was determined from that *sam* file since there was only one mapping for each read. Only the *sam* lines with FLAG values of 0 or 16 were considered for analysis.

### Supplementary Material 3 - Database creation

The databases were prepared by downloading the sequences from the refseq database and separating the genome sequences in files where every file contains sequences from one species. The taxonomy files, *nodes.dmp* and *names.dmp* were also downloaded alongside the genome sequences.

For the tools that use protein sequences the different protein database was downloaded, also from the refseq database, with different *nodes.dmp* and *names.dmp* files.

The Kraken's database was built by adding the string *kraken:taxid|XXX* to the sequence ID, where the *XXX* represents the desired taxon ID. Each fasta file was added to the kraken's database library with the command: *kraken2-build --add-to-library chr1.fa --db $DBNAME* Finally, the database index was built with the command: *kraken2-build --build --db $DBNAME*

For the centrifuge database the mapping file *seqid2taxid.map*, that contains mapping of sequence IDs to the tax IDs was created. Also the *input-sequences.fna* was created by concatenating all the fasta files of the genome sequences. The database index was built with the command: *centrifuge-build -p 4 --conversion-table seqid2taxid.map --taxonomy-tree nodes.dmp --name-table names.dmp input-sequences.fna database_name*

The Clark's database was built by creating the *Custom* directory inside *DIR_DB* directory and placing all the genome fasta sequences inside. The database index was created with the command: *./set_targets.sh DIR_DB custom*

The discriminative spaced k-mers for Clark S were built on top of this database by running the command *./buildSpacedDB.sh*.

The file containing concatenated genome sequences was used for MetaMaps's database index creation, with the command: *perl buildDB.pl --DB database_name --FASTAs all.fna --taxonomy download/taxonomy_uniqueIDs*

MEGAN-LR uses Last aligner for the mapping part of the pipeline. For custom database containing nucleotide sequences, LAST index was created using the following instruction:

*lastdb -cR01 last_custom_idx NT.fa*

It should be noted that creating a last index took 1-2 days on the server we used. Also, indexing nucleotide sequences (NCBI bacteria, archaea, and human genomes), the index created was quite large, taking around 500GB of disk space.

For the minimap2 and Ram all the concatenated fasta files of the genome sequences were passed to the tools as the target sequence parameter. For these two tools, a preprocessed database index was not created since the tools don't require one to be built.

Kaiju's database index was created from the protein database file with modified sequence headers in the format ">orderNumber_taxID". Then the index was created with the following commands:

*kaiju-mkbwt -n 24 -a ACDEFGHIKLMNPQRSTVWY -o database_index_filename database_filename*

*kaiju-mkfmi database_index_filename*

deSAMBA database index was created from the database file with the modified sequence headers in the format ">tid|tax_id|". The index was built with the following command:
*bash build-index <database_filename> <database_index_name>*

### Supplementary Figure 1

**Read level classification F1 macro average,** c**omparison between species and genus level classification.** Classification for each read is compared to the ground truth and used to calculate F1 value for each organism in the sample separately. Calculated F1 values are then averaged over all organisms to obtain F1 macro average. Plot a) shows species level classification for which reads are considered correctly classified if classified to a correct species. Plot b) shows genus level classification for which reads are considered correctly classified if classified to a correct genus. Results for MEGAN are unavailable for PB3 dataset.

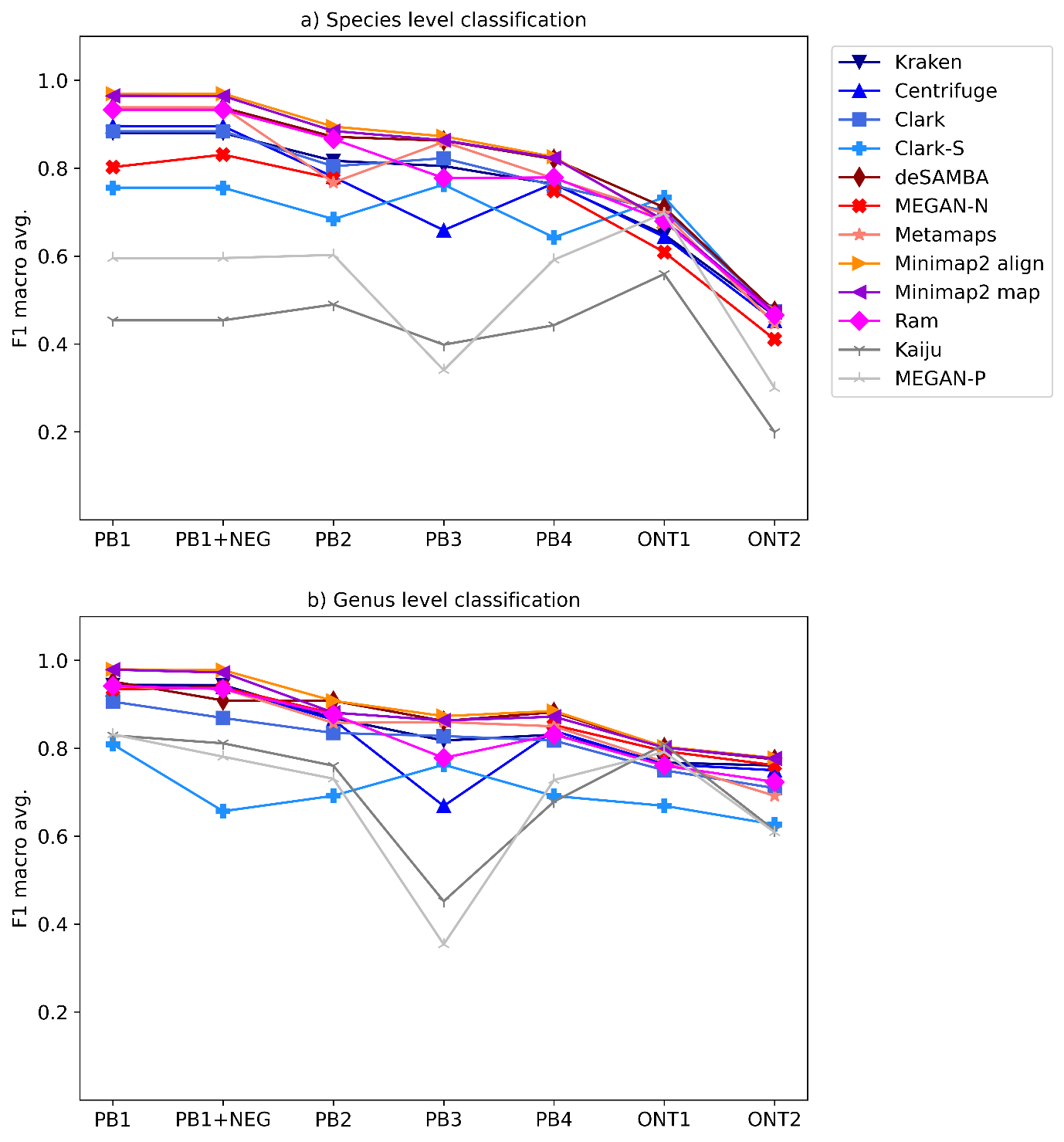

### Supplementary Figure 2

**In-sample abundance error boxplots for all datasets.** Abundance estimation error is calculated by comparing abundances calculated for each tool to the ground truth (L1 distance). Errors are calculated separately for each organism in the sample. The results are plotted as a box-plot. Data for MEGAN-N are unavailable for datasets PB3, PB_ATCC and PB_Zymo. Since PB3 has only 2 species,plotting 2 points instead of a full boxplot. Outliers (Supplementary Table 11) that appear on the same level for all tools represent species not present in the database (most noticeable on datasets ONT1 and PB_Zymo).

| 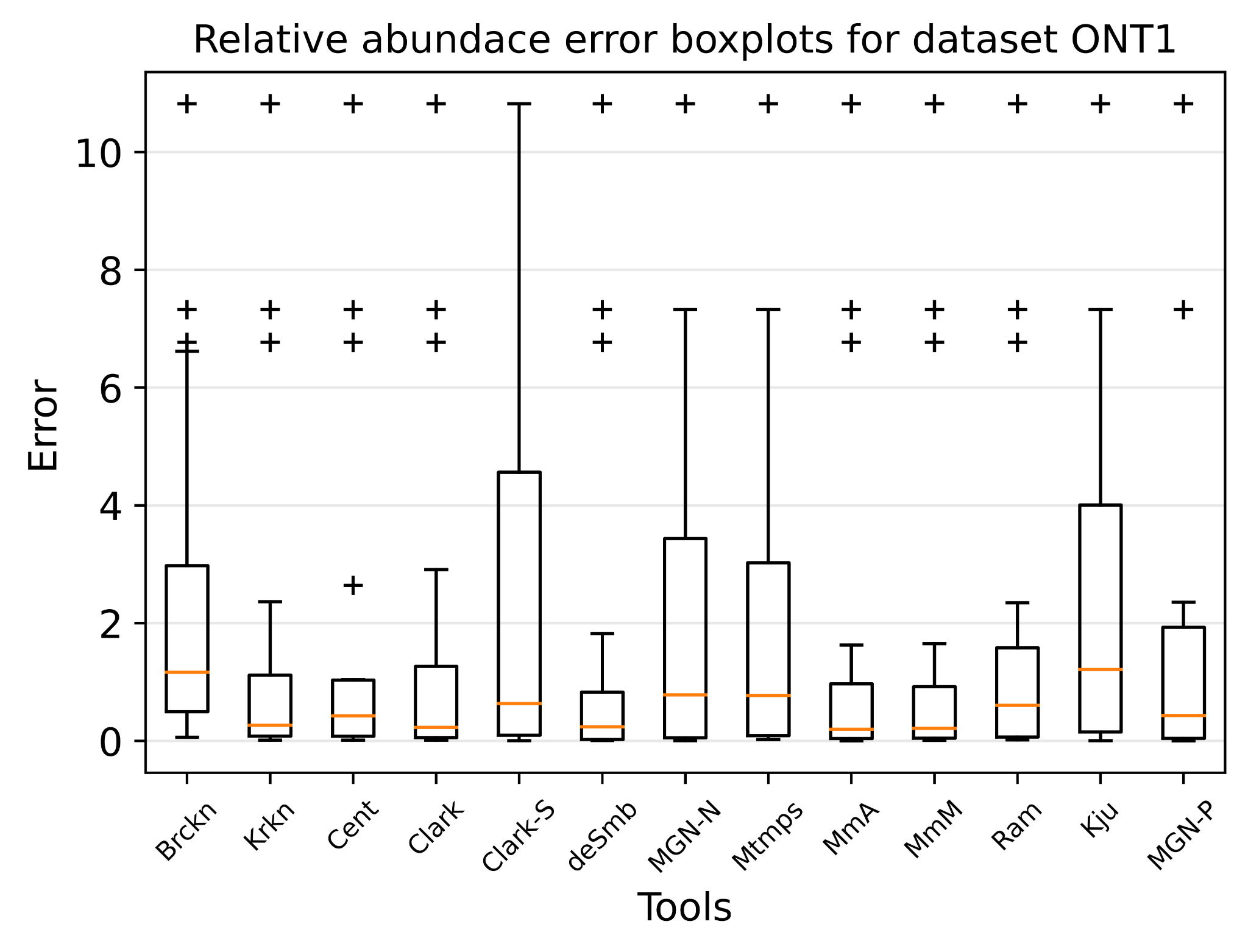 | 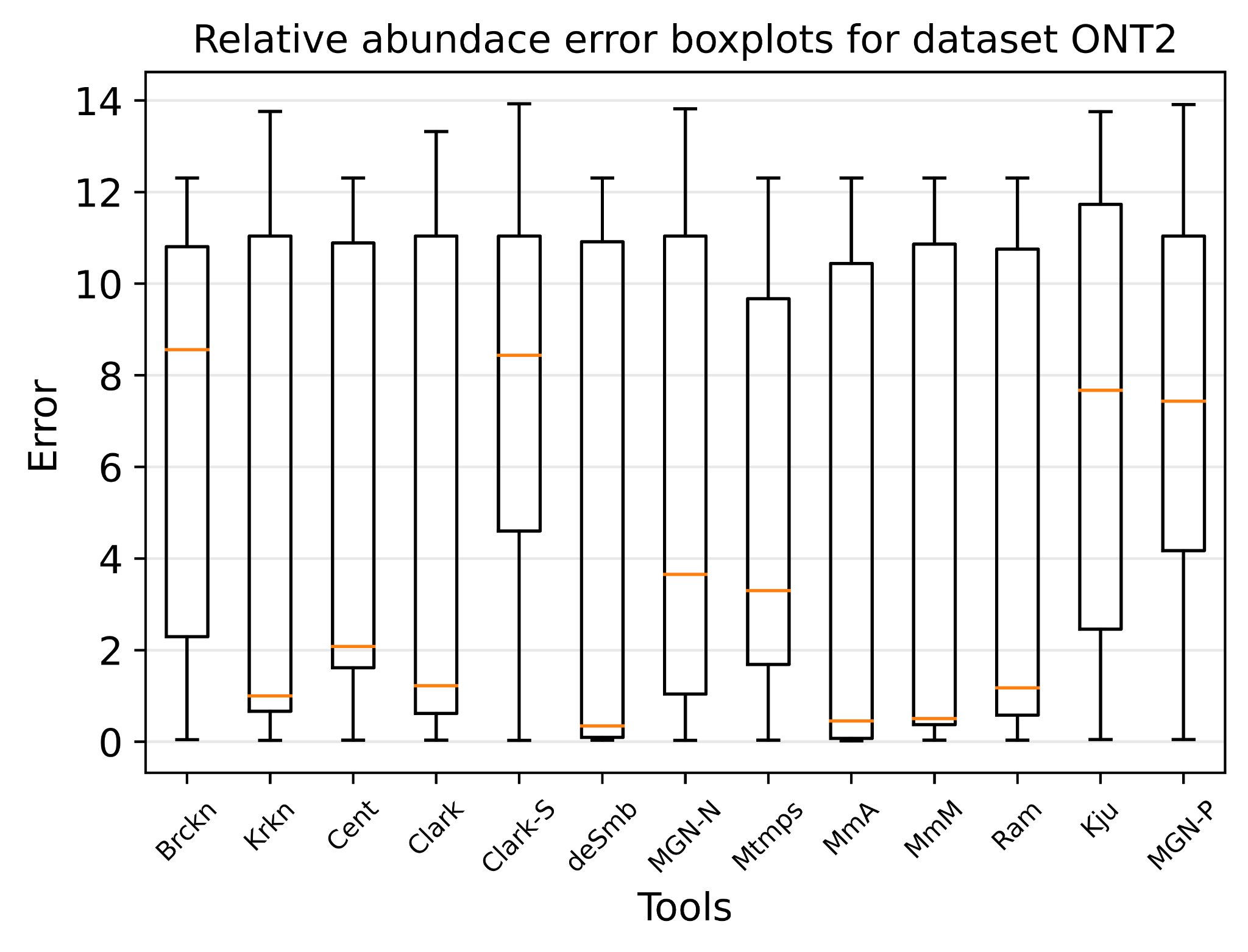 |
| --- | --- |
| 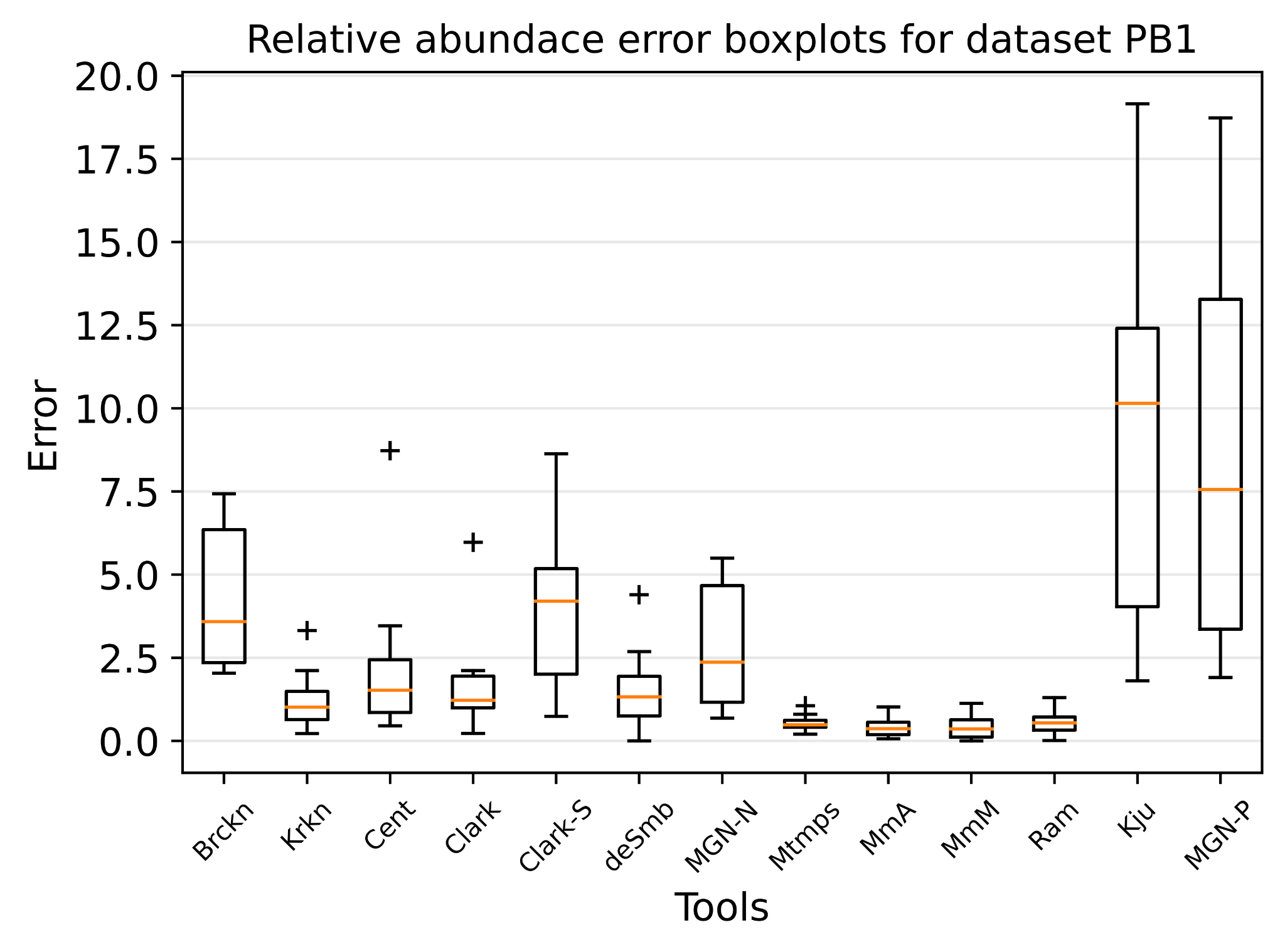 | 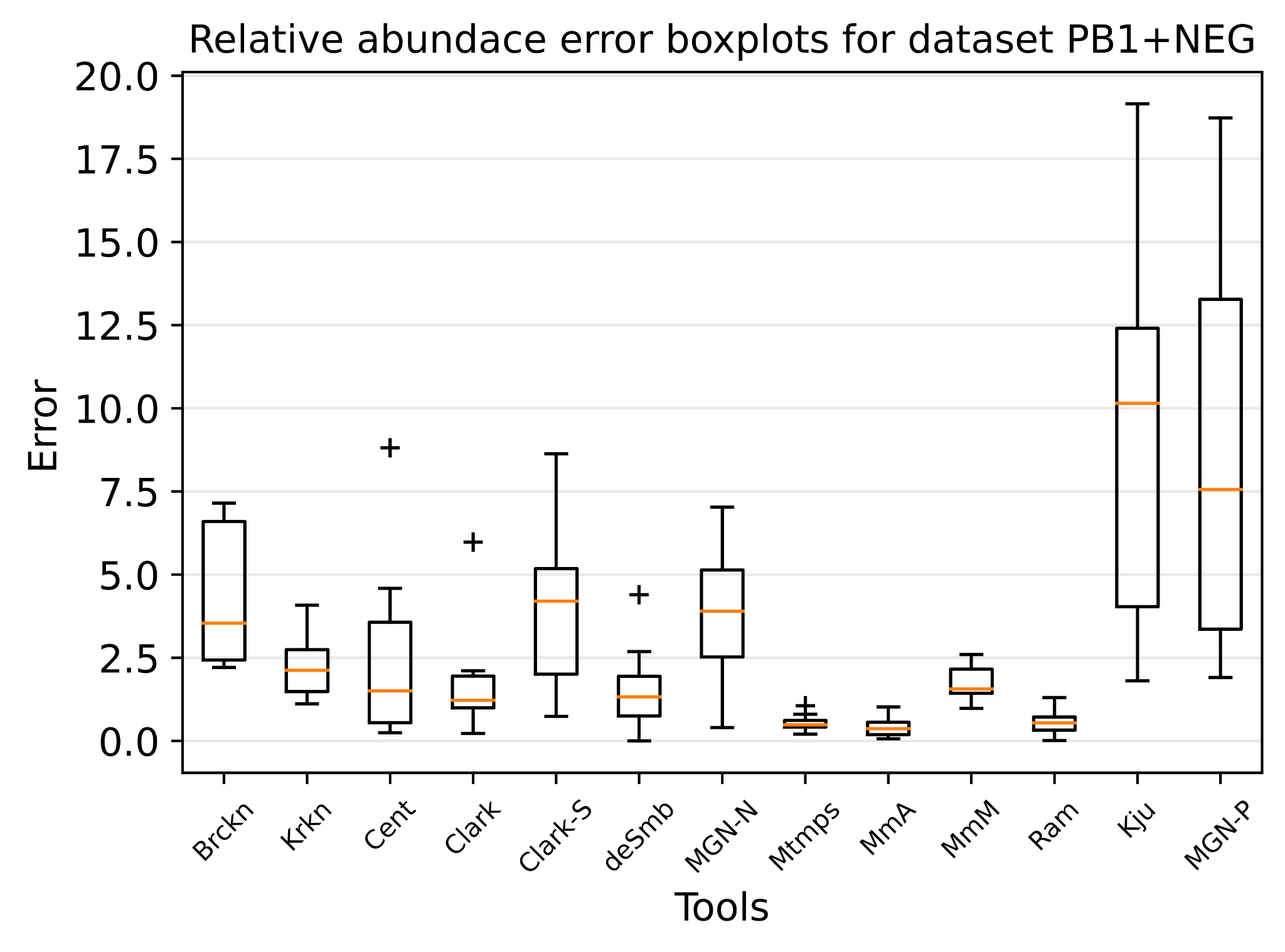 |
| 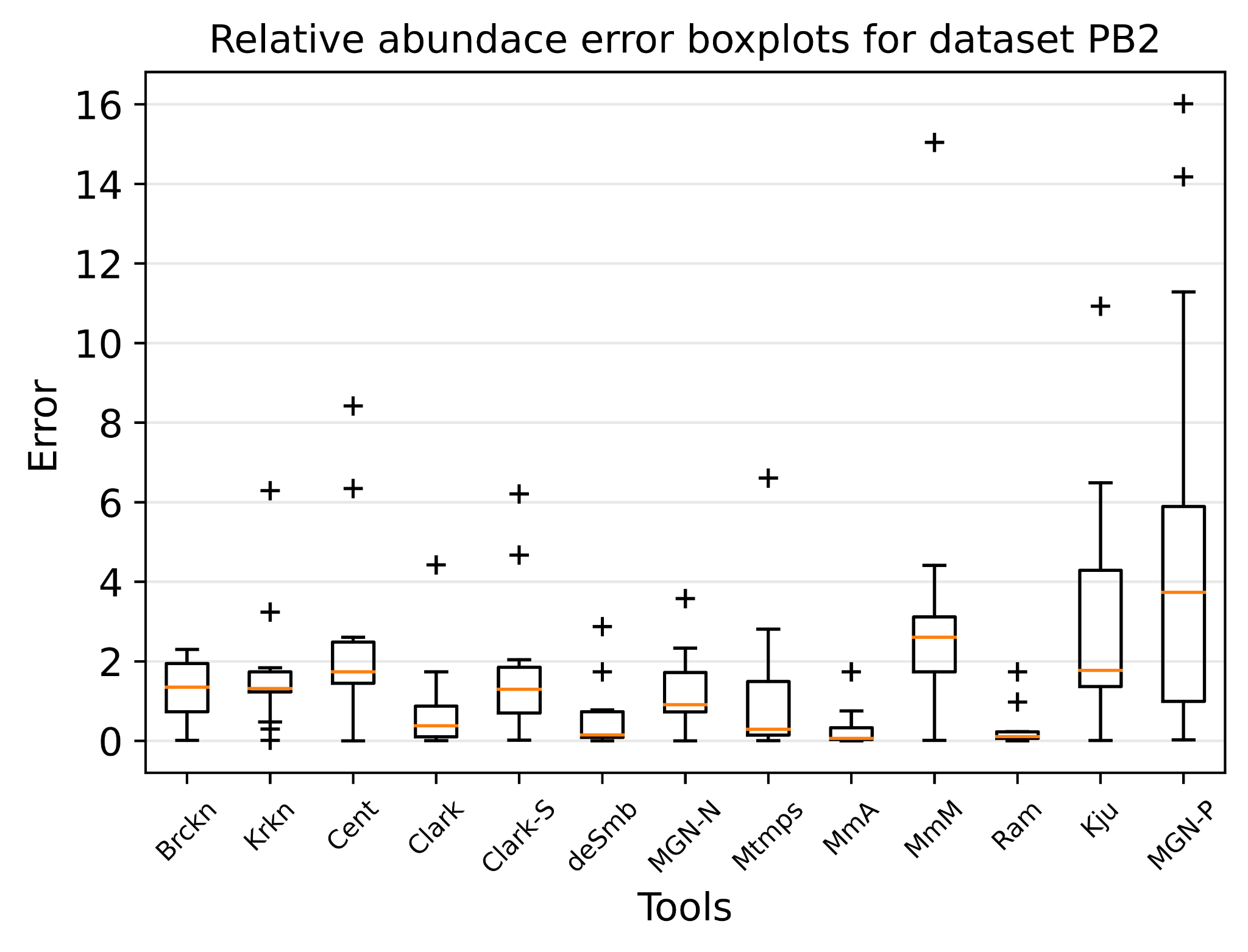 | 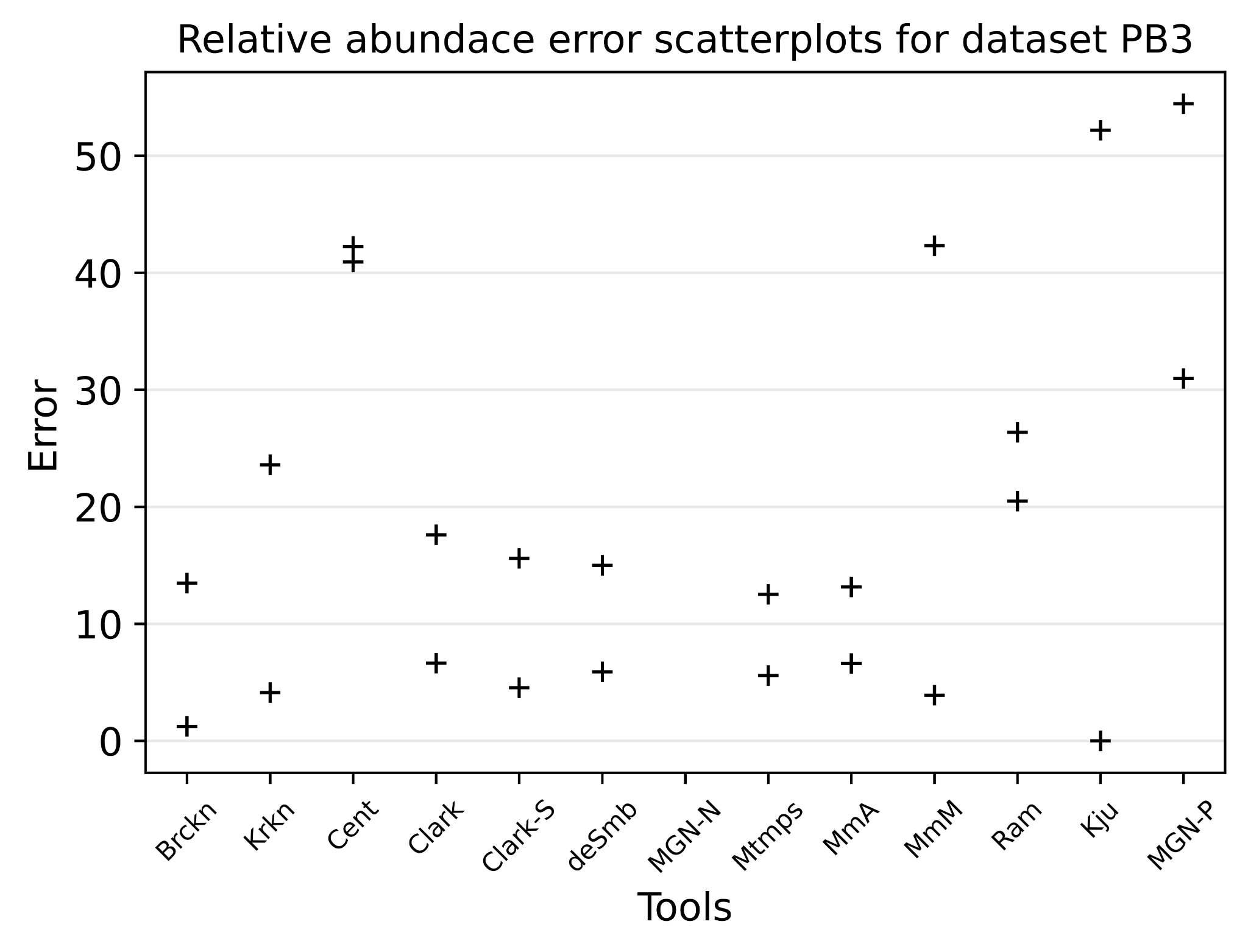 |
| 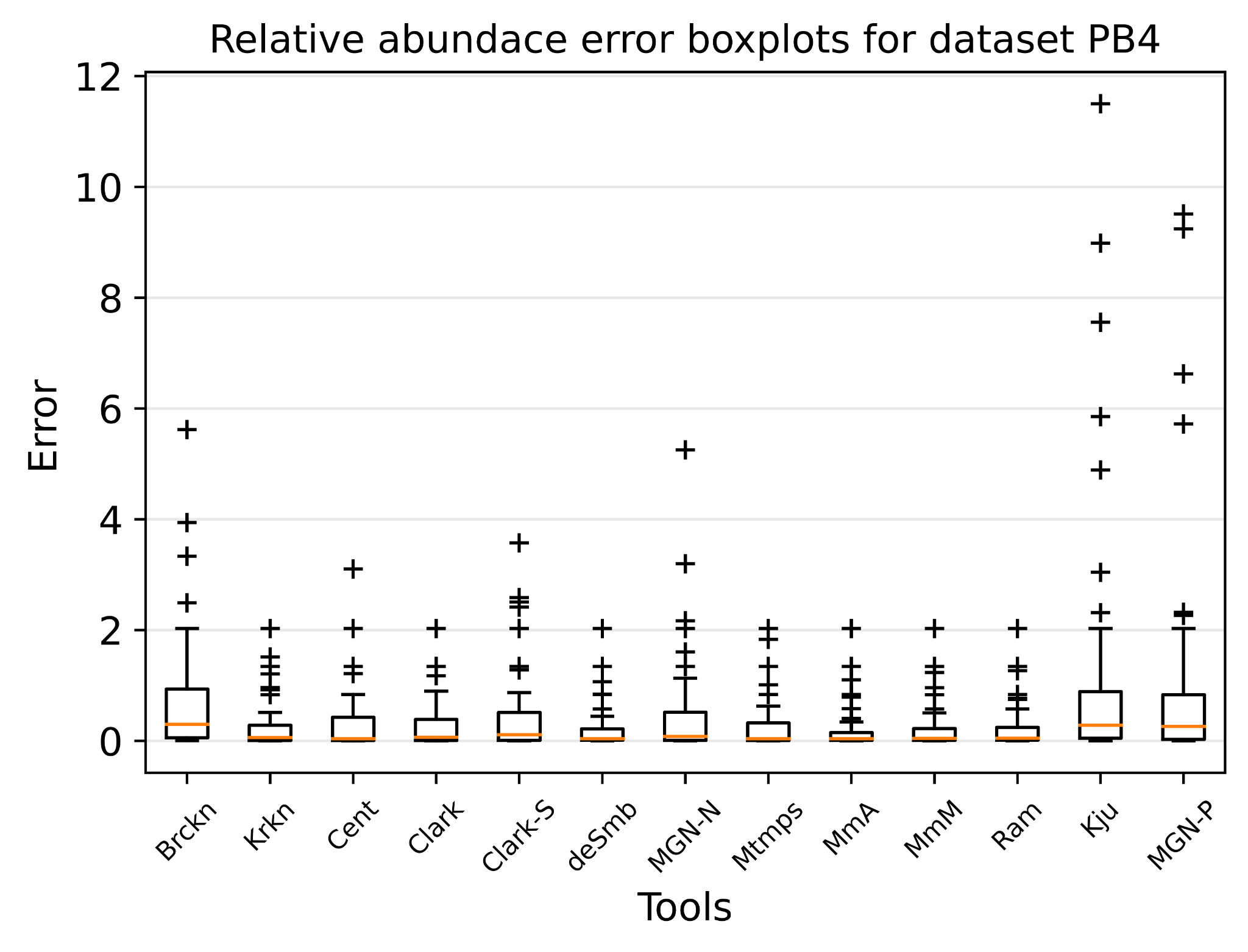 | 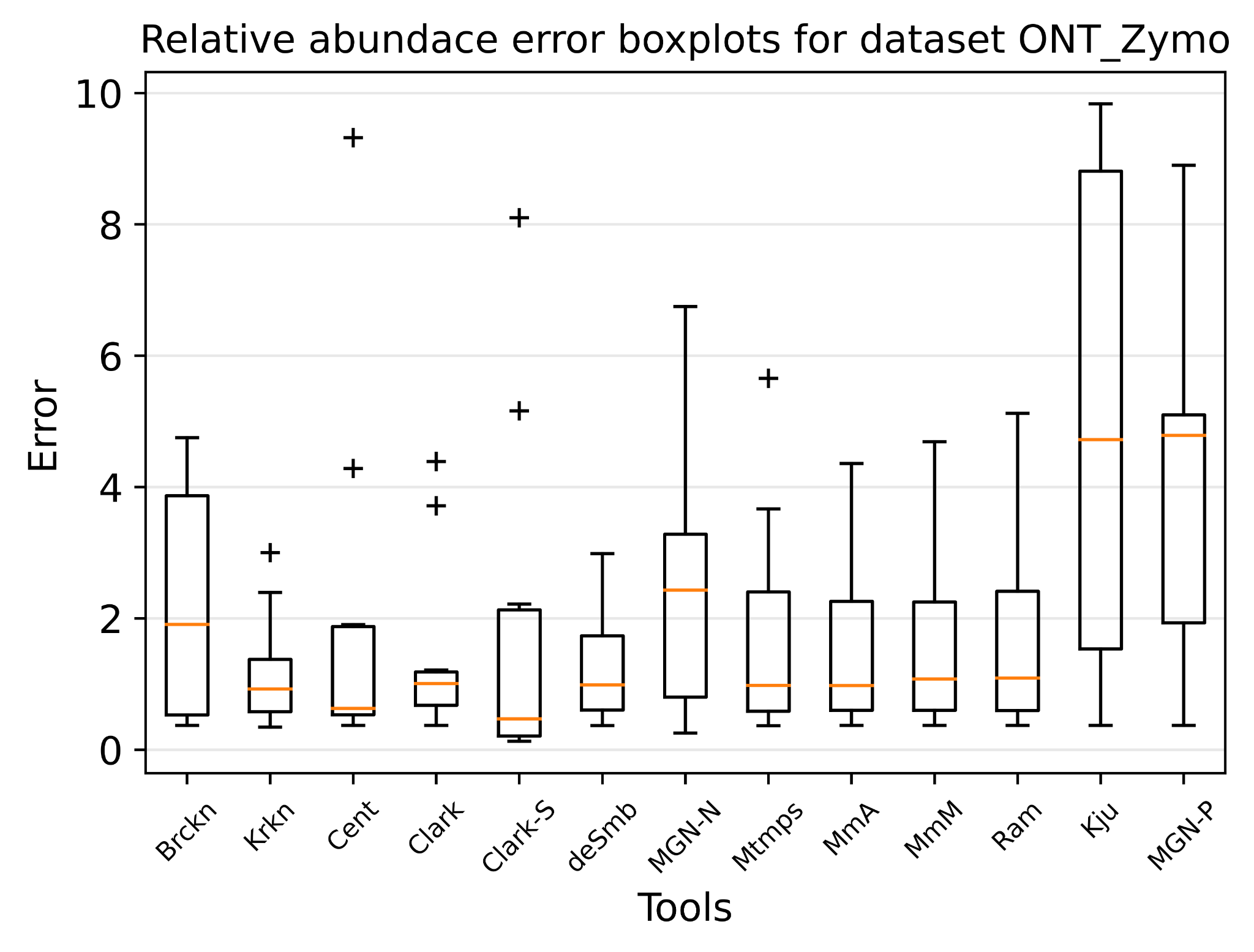 |
| 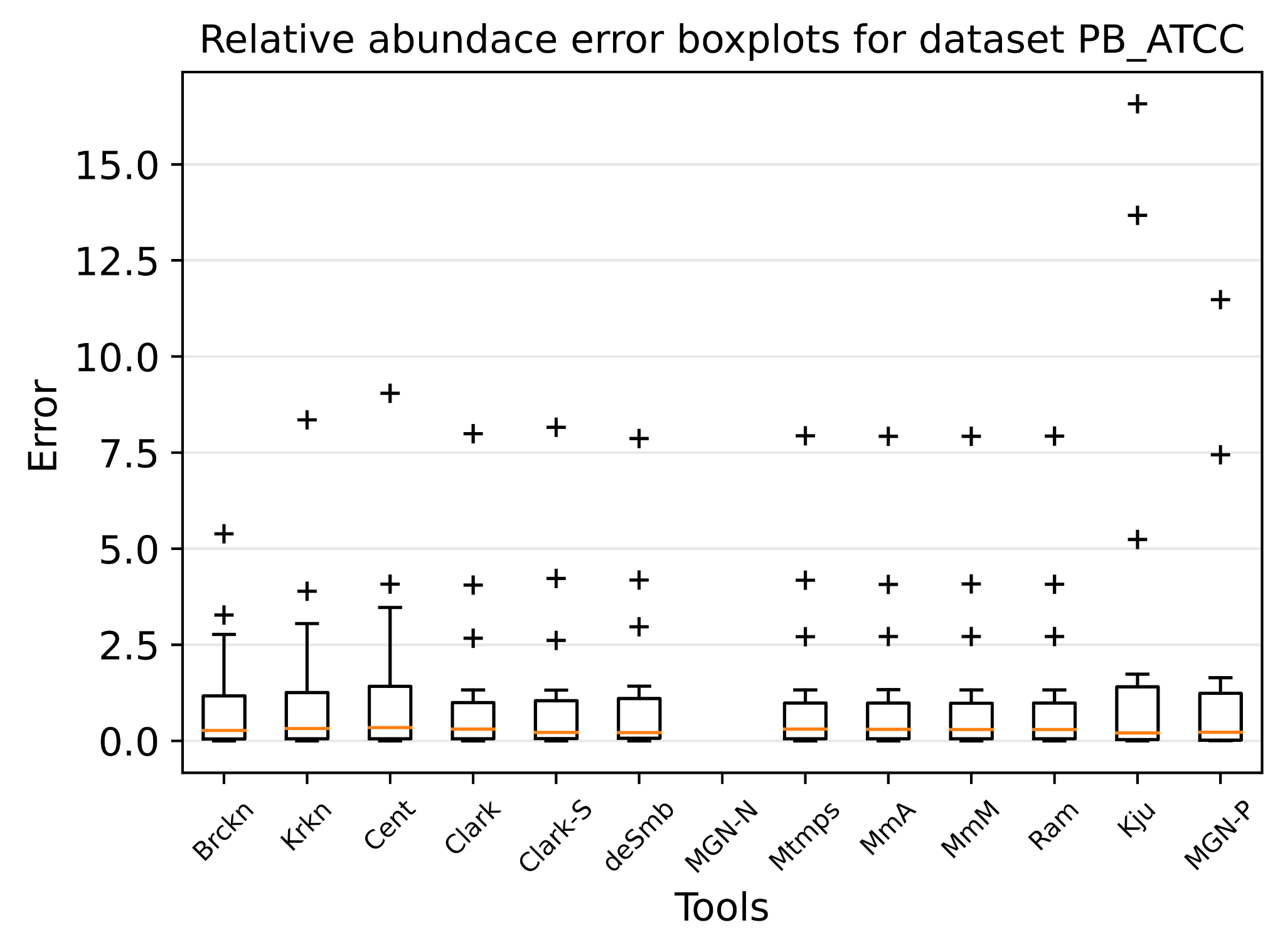 | 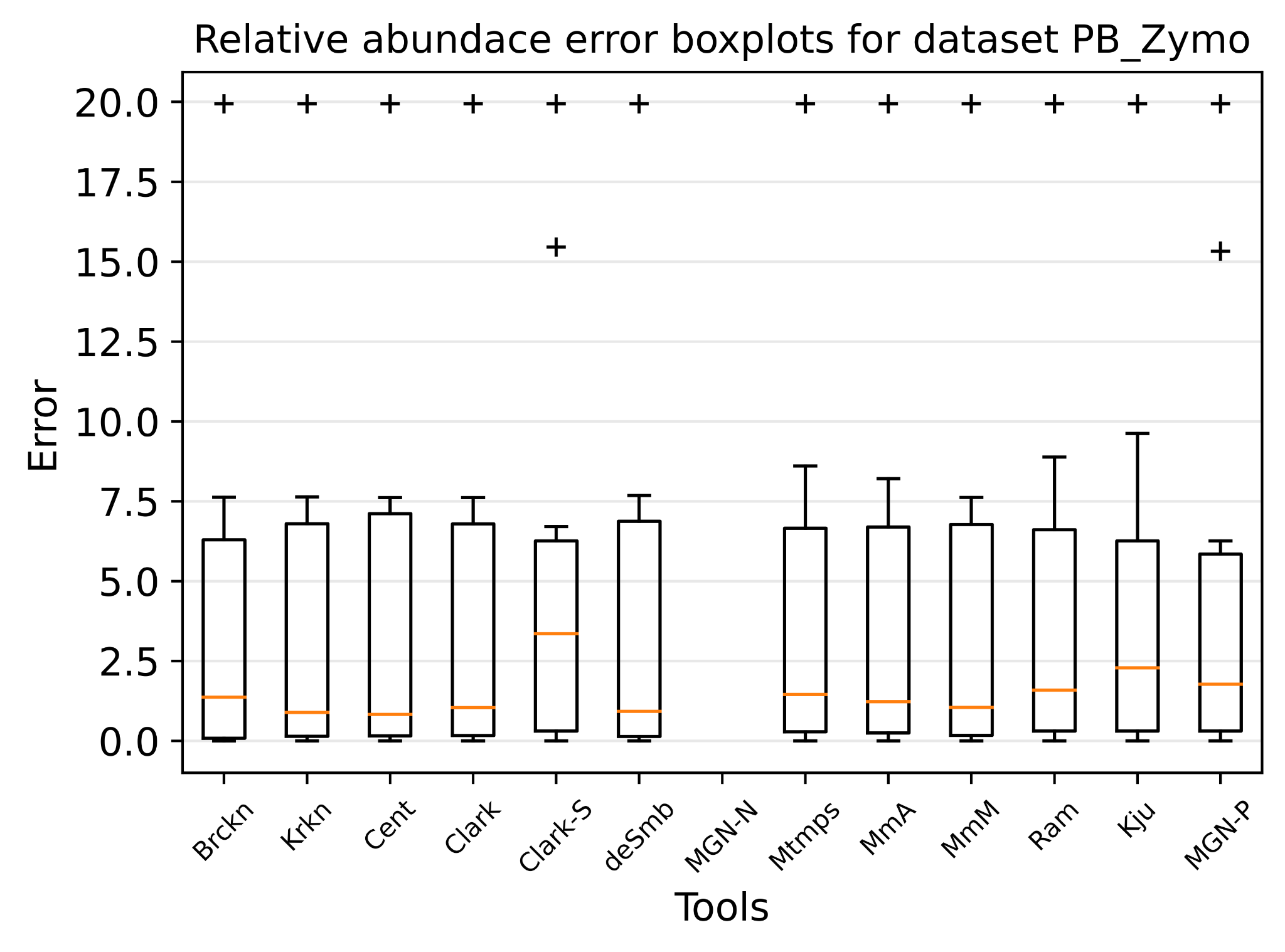 |

### Supplementary Figure 3

**In-sample abundance error boxplots for mock community datasets - comparison between read counts and sum of reads’ lengths.** Abundance estimation error is calculated by comparing abundances calculated for each tool to the ground truth (L1 distance). Errors are calculated separately for each organism in the sample. The plots are shown for two definitions of relative abundance: 1) percentage of reads classified to a species based on read counts, 2) percentage of total read length of reads classified to a species. The results are plotted as a box-plot. Data for MEGAN-N are unavailable for datasets PB_ATCC and PB_Zymo due to long running time. Moreover, It is important to note that Bracken produces only read counts assigned to a taxonomic rank.

| 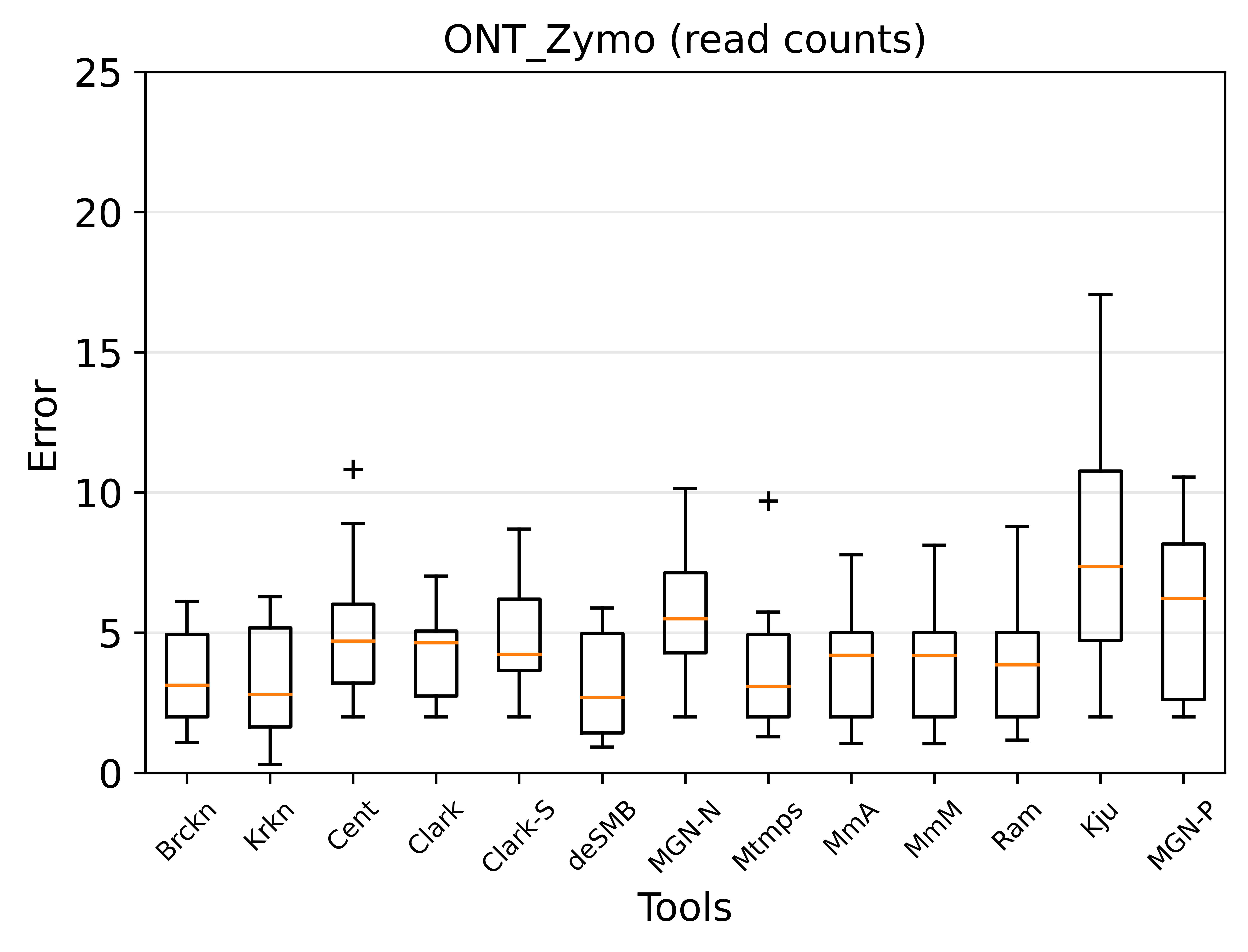 | 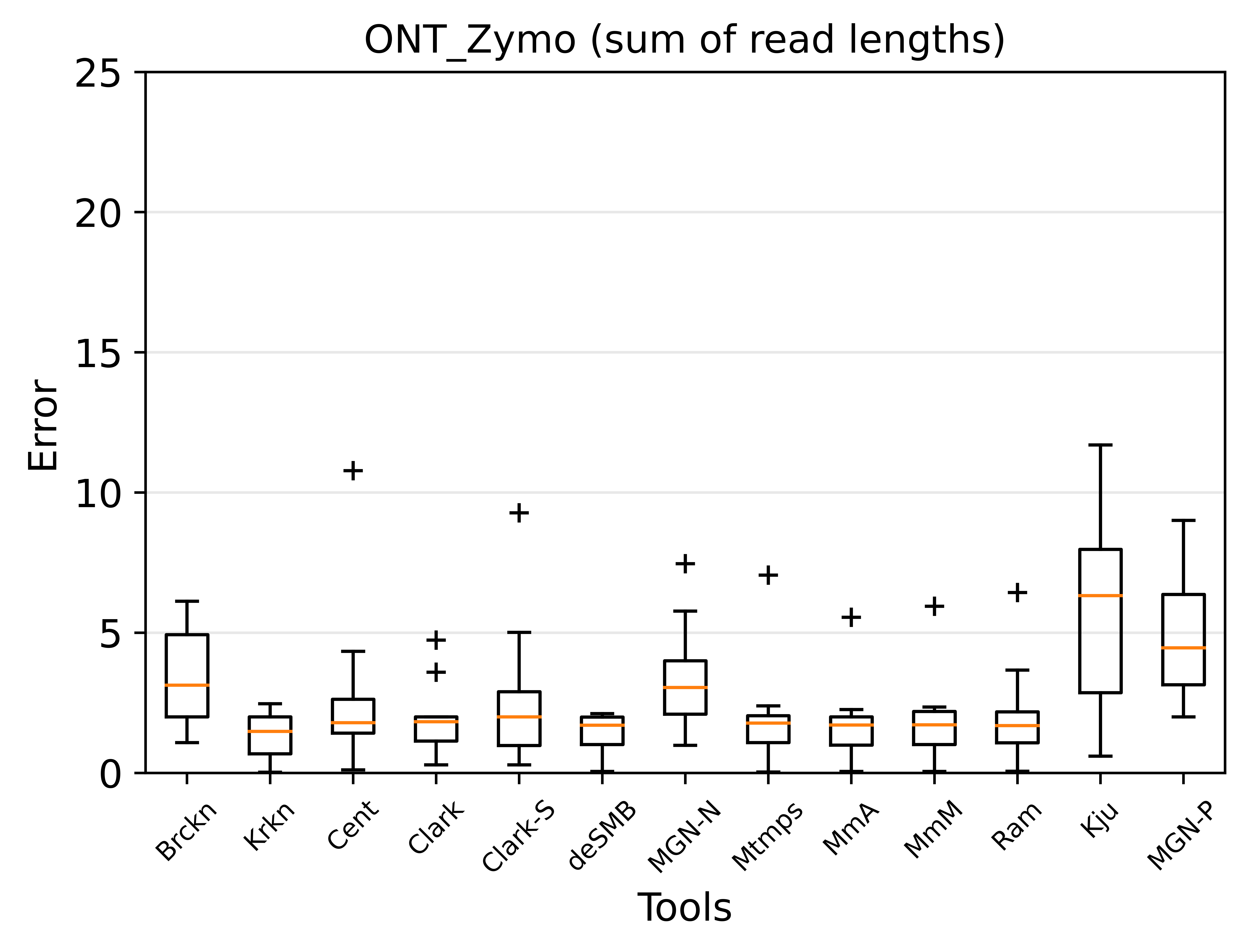 |
| --- | --- |
| 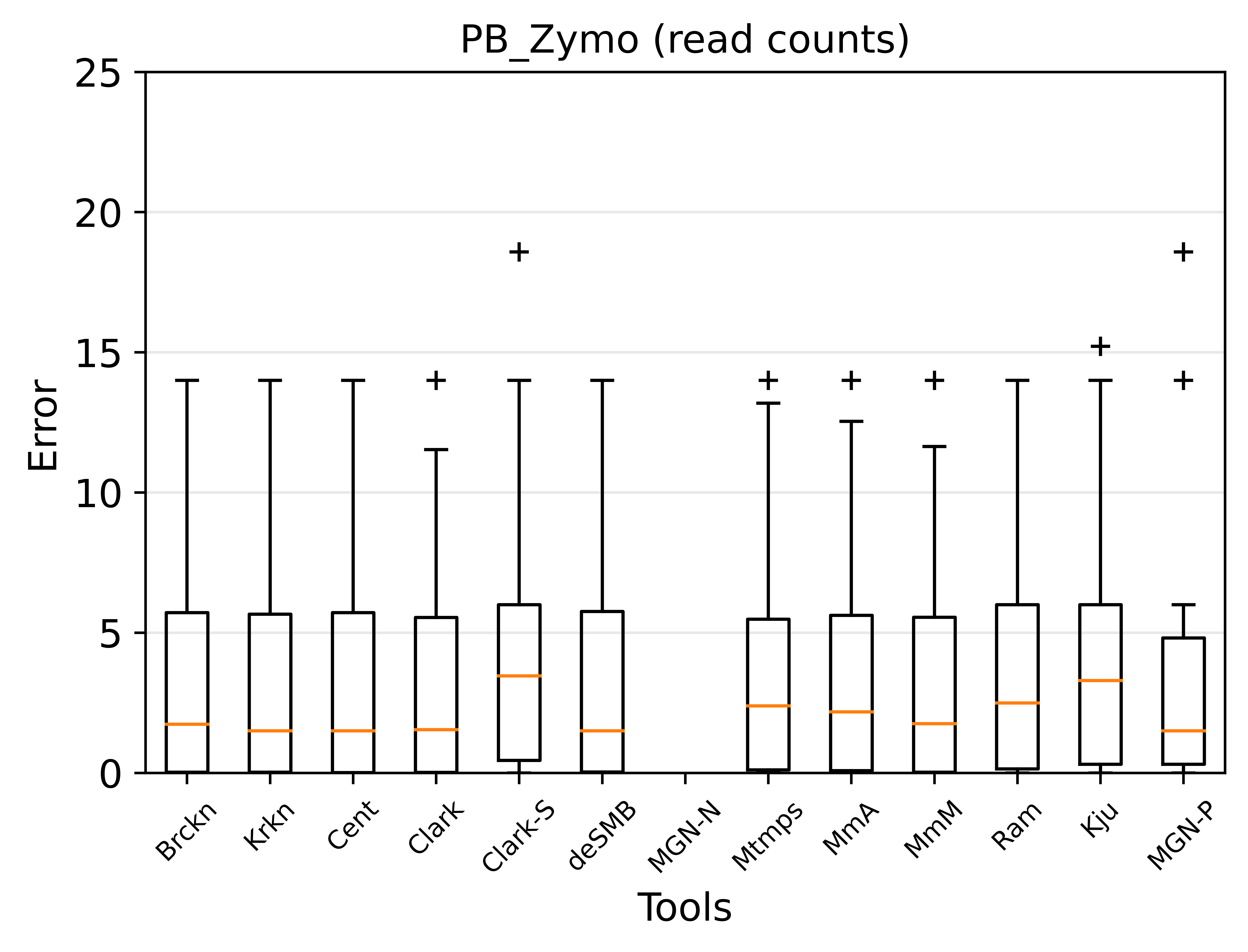 | 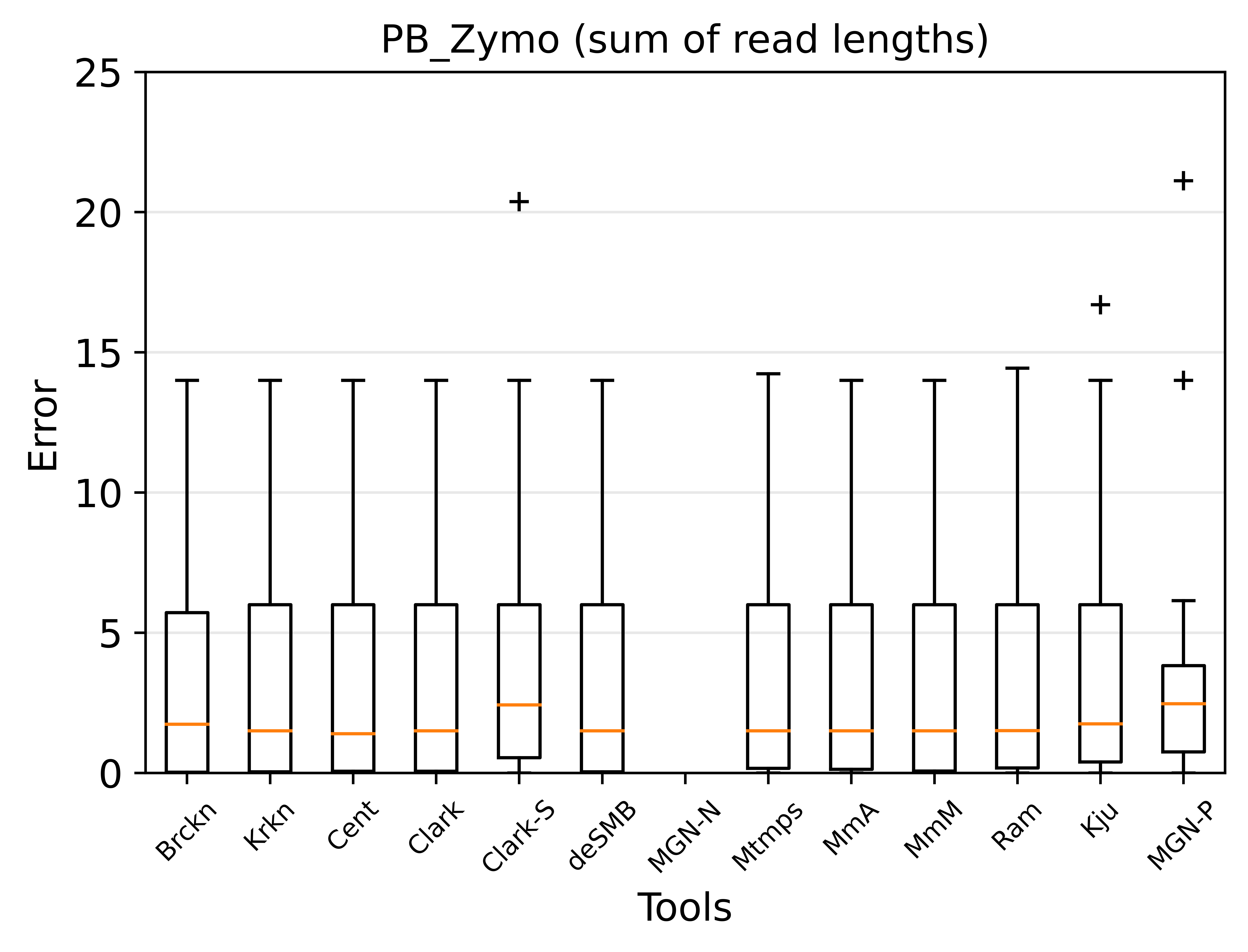 |
| 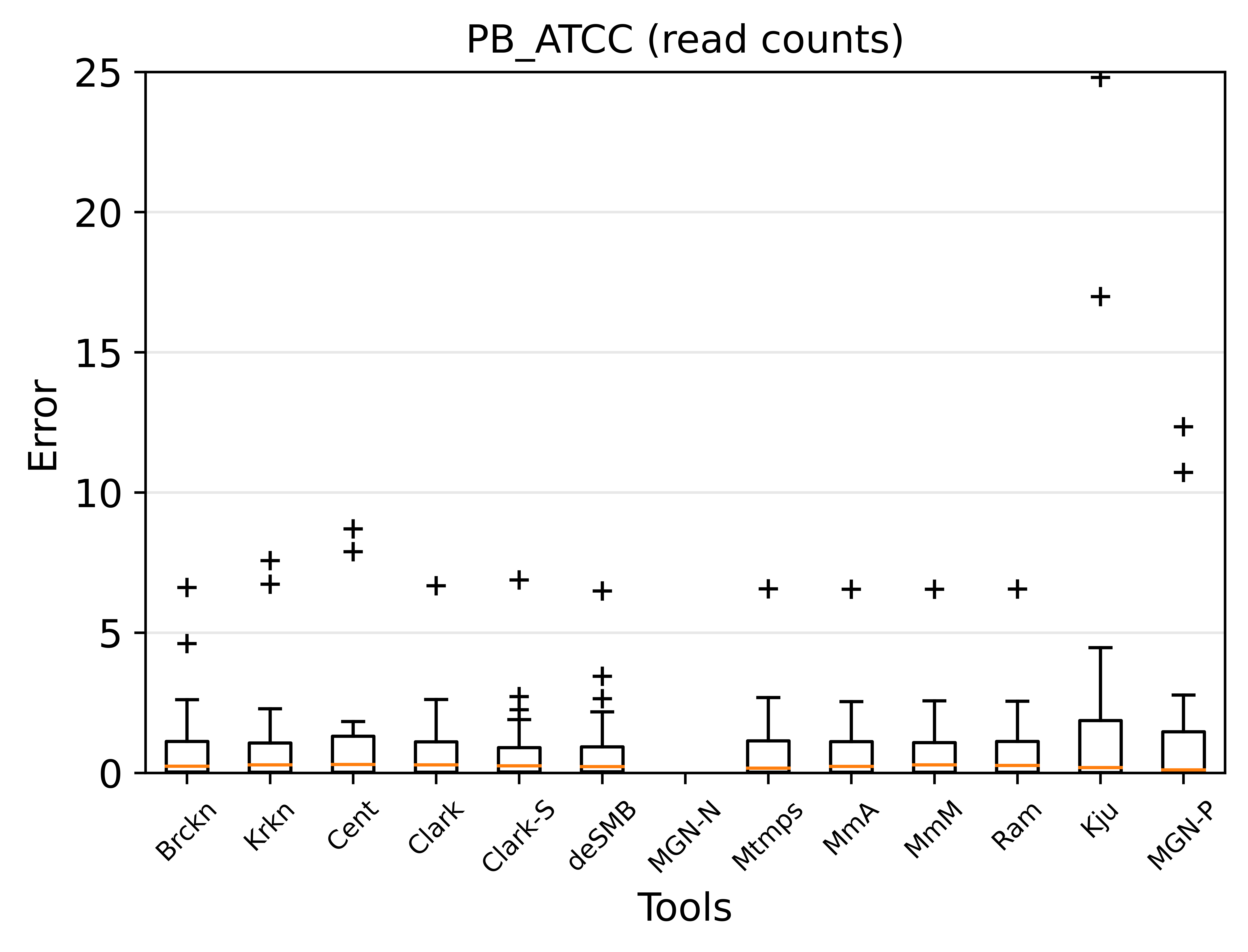 | 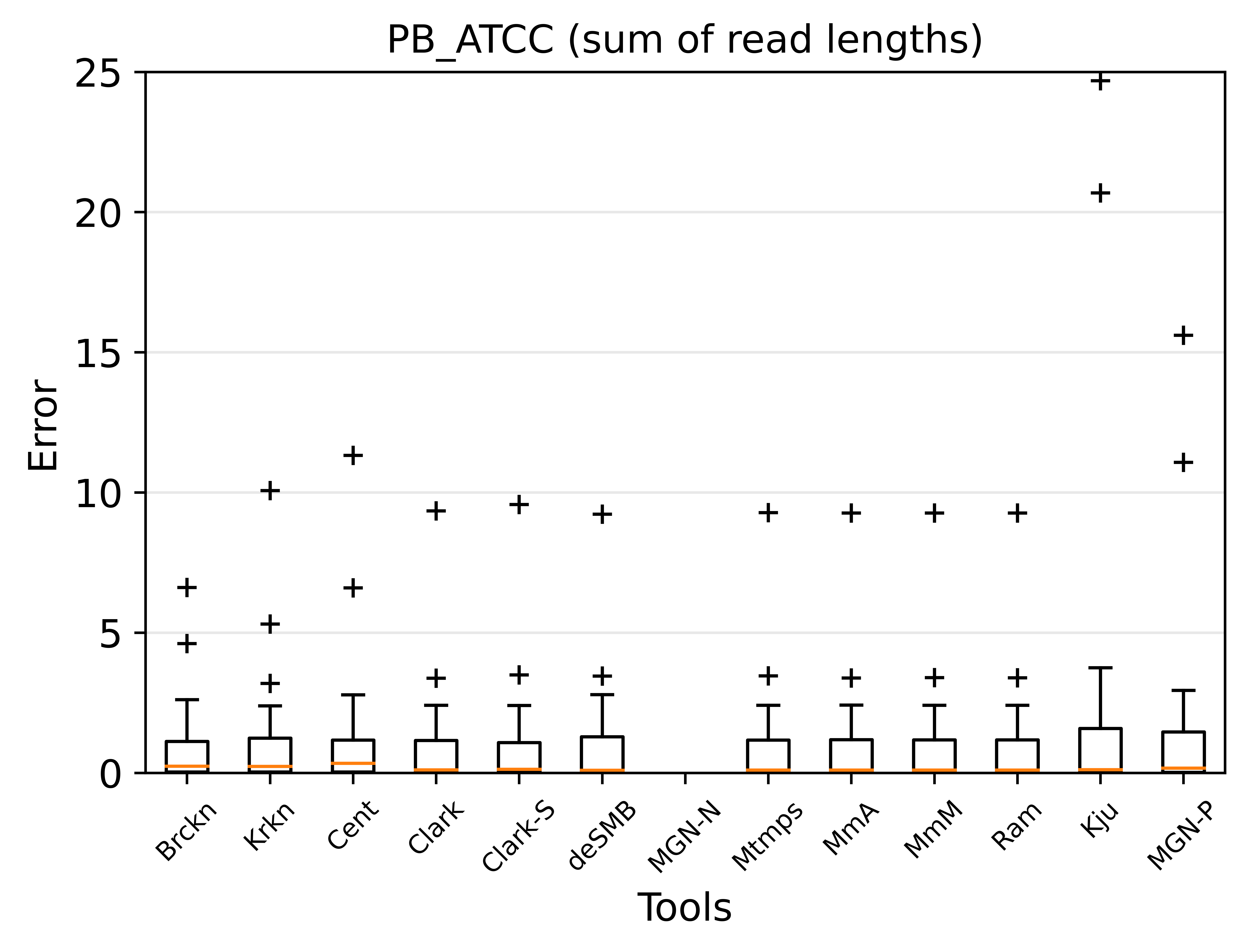 |

### Supplementary Table 1

**Total absolute abundance estimation error using 30% of longest reads and all reads.**

The table shows the total abundance estimation error for real datasets for two different cases. 1) Using only 30% of the longest reads, 2) using all reads. The error is calculated by calculating the absolute value of the difference between abundance reported by a tool and true abundance and summing it up across all present and reported organisms. Results for MEGAN on the nucleotide database are unavailable for datasets PB_Zymo and PB_ATCC. The ONT_Zymo dataset consists of 10 different species obtained by the GridION sequencer. The PB_ATCC dataset consists of 20 different species sequenced using PacBio HiFi technology. The PB_Zymo dataset consists of 17 different species sequenced using PacBio HiFi technology.

| **Data set** | **Reads** | **Kraken2** | **Centrifuge** | **CLARK** | **CLARK-S** | **Metamaps** | **MEGAN-N** | **deSAMBA** | **Minimap2 align** | **Minimap2 map** | **Ram** | **Kaiju** | **MEGAN -P** |
| --- | --- | --- | --- | --- | --- | --- | --- | --- | --- | --- | --- | --- | --- |
| **ONT Zymo** | **Long reads** | 26.8 | 32.4 | 24.7 | 31.3 | 24.2 | 26.3 | 29.9 | 24.1 | 24.2 | 24,2 | 51.4 | 38.8 |
|  | **All reads** | 18.9 | 26.6 | 19.5 | 19.2 | 17.5 | 27.2 | 20.3 | 16.5 | 16.9 | 18.6 | 59.0 | 41.6 |
| **PB ATCC** | **Long reads** | 37.9 | 38.3 | 37.3 | 37.5 | 37.3 | - | 37.8 | 37.6 | 37.6 | 37.5 | 56.7 | 44.7 |
|  | **All reads** | 25.8 | 26.1 | 21.4 | 21.1 | 21.1 | - | 23.4 | 21.4 | 21.6 | 21.5 | 44.8 | 29.2 |
| **PB Zymo** | **Long reads** | 124.7 | 125.4 | 124.4 | 103.1 | 122.6 | - | 125.3 | 123.5 | 124.7 | 122.8 | 128.9 | 95.0 |
|  | **All reads** | 110.1 | 111.0 | 109.7 | 93.1 | 108.6 | - | 110.7 | 109.1 | 109.9 | 109.6 | 115.7 | 83.5 |

### Supplementary Table 2

**Resource usage.** The table shows running time (in seconds) and memory usage (in GB) for all tools and all synthetic and mock community datasets (all roughly 1Gbp in size).

| **Execution time / s** | | | | | | | | | | | |  |  |
| --- | --- | --- | --- | --- | --- | --- | --- | --- | --- | --- | --- | --- | --- |
| **Dataset**  **/ tool** | **Kraken2** | **Bracken** | **Centrifuge** | **CLARK** | **CLARK-S** | **Metamaps** | **MEGAN-N** | **deSAMBA** | **Minimap2 align** | **Minimap2 map** | **Ram** | **Kaiju** | **MEGAN -P** |
| **ONT1** | 314 | 126166 | **275** | 974 | 3913 | 37839 | 67090 | 895 | 3145 | 1710 | 1482 | 525 | 21371 |
| **ONT2** | 315 | 147207 | **291** | 954 | 3917 | 39844 | 75214 | 988 | 2874 | 1808 | 1509 | 528 | 16183 |
| **PB1** | 312 | 137017 | **284** | 972 | 3942 | 54829 | 84852 | 1337 | 3797 | 1890 | 1541 | 592 | 24971 |
| **PB1+NEG** | 321 | 145769 | **296** | 968 | 4100 | 68941 | 157526 | 1235 | 3778 | 1799 | 1600 | 600 | 29149 |
| **PB2** | 326 | 191661 | **296** | 993 | 4114 | 58815 | 119849 | 955 | 2641 | 1647 | 1607 | 605 | 31099 |
| **PB3** | 320 | 224048 | **309** | 979 | 4075 | 145416 | - | 901 | 1904 | 1541 | 1591 | 610 | 23254 |
| **PB4** | 308 | 98826 | **267** | 963 | 3862 | 50180 | 58238 | 996 | 2843 | 1597 | 1511 | 526 | 19252 |
| **ONT_Zymo** | 327 | 144387 | **305** | 979 | 4084 | 70622 | 160225 | 1344 | 4220 | 2283 | 1697 | 341 | 32013 |
| **PB_ATCC** | **317** | 68971 | 329 | 975 | 3957 | 76897 | - | 1400 | 3044 | 1928 | 1303 | 347 | 33430 |
| **PB_Zymo** | 317 | 179206 | **292** | 953 | 3996 | 63142 | - | 1318 | 2364 | 1604 | 1272 | 329 | 32508 |
| **Memory / GB** | | | | | | | | | | | |  |  |
| **Dataset**  **/ tool** | **Kraken2** | **Bracken** | **Centrifuge** | **CLARK** | **CLARK-S** | **Metamaps** | **MEGAN**  **-N** | **deSAMBA** | **Minimap2 align** | **Minimap2 map** | **Ram** | **Kaiju** | **MEGAN -P** |
| **ONT1** | 43.04 | 45.31 | 37.00 | 119.56 | 271.24 | 205.67 | 26.87 | 154.48 | 39.82 | 34.18 | **14.05** | 55.28 | 93.29 |
| **ONT2** | 43.12 | 25.38 | 36.91 | 119.39 | 271.46 | 208.46 | 78.91 | 161.34 | 47.40 | 31.34 | **14.06** | 55.41 | 54.81 |
| **PB1** | 43.03 | 25.47 | 37.08 | 118.91 | 271.16 | 208.46 | 30.43 | 159.28 | 28.60 | 19.98 | **14.10** | 55.28 | 55.75 |
| **PB1+NEG** | 43.01 | 25.39 | 37.02 | 119.15 | 271.17 | 208.46 | 30.94 | 159.32 | 27.52 | 21.04 | **14.20** | 55.28 | 55.33 |
| **PB2** | 43.04 | 24.39 | 36.56 | 120.29 | 271.42 | 146.29 | 108.22 | 154.63 | 31.30 | 22.35 | **13.97** | 55.28 | 54.72 |
| **PB3** | 42.99 | 25.38 | 36.09 | 120.61 | 271.32 | 208.45 | - | 143.55 | 24.02 | 21.47 | **14.26** | 55.28 | 22.73 |
| **PB4** | 43.04 | 25.42 | 36.67 | 119.33 | 271.25 | 208.46 | 29.90 | 161.49 | 27.32 | 22.71 | **14.13** | 55.28 | 87.45 |
| **ONT_Zymo** | 43.02 | 25.41 | 37.06 | 120.08 | 271.14 | 208.46 | 42.57 | 179.14 | 41.56 | 38.23 | **14.27** | 55.28 | 85.34 |
| **PB_ATCC** | 43.01 | 24.37 | 36.00 | 120.05 | 271.25 | 208.46 | - | 213.95 | 28.77 | 19.77 | **9.07** | 55.28 | 87.10 |
| **PB_Zymo** | 42.98 | 24.42 | 35.94 | 120.06 | 271.22 | 208.41 | - | 189.01 | 26.19 | 21.00 | **9.15** | 55.28 | 77.46 |

### Supplementary Table 3

**Percentage of reads classified for all datasets and tools.**

| **Dataset**  **/ tool** | **Number of reads** | **Kraken2** | **Centrifuge** | **CLARK** | **CLARK-S** | **Metamaps** | **MEGAN-N** | **deSAMBA** | **Minimap2 align** | **Minimap2 map** | **Ram** | **Kaiju** | **MEGAN -P** |
| --- | --- | --- | --- | --- | --- | --- | --- | --- | --- | --- | --- | --- | --- |
| **ONT1** | 100k | 85.6 | 84.6 | 79.6 | 57.3 | 76.3 | 76.5 | 87.8 | 88.0 | 88.4 | 74.2 | 58.5 | 70.0 |
| **ONT2** | 74k | 82.2 | 84.1 | 74.9 | 55.2 | 68.6 | 70.0 | 85.6 | 86.1 | 86.4 | 72.1 | 43.9 | 50.5 |
| **PB1** | 100k | 86.3 | 80.5 | 81.9 | 61.0 | 89.7 | 74.3 | 88.1 | 96.5 | 97.0 | 90.3 | 34.5 | 42.8 |
| **PB1+NEG** | 120k | 79.1 | 74.6 | 68.5 | 51.0 | 75.2 | 58.8 | 73.7 | 80.7 | 87.9 | 75.5 | 28.9 | 35.8 |
| **PB2** | 100k | 85.5 | 82.4 | 73.8 | 65.1 | 75.8 | 70.6 | 79.6 | 79.3 | 94.1 | 76.2 | 50.9 | 55.4 |
| **PB3** | 100k | 99.7 | 99.7 | 99.4 | 99.5 | 99.8 | - | 99.8 | 99.1 | 100 | 98.7 | 94.0 | 57.2 |
| **PB4** | 100k | 89.3 | 89.4 | 86.4 | 67.7 | 92.0 | 76.2 | 97.1 | 97.2 | 98.3 | 90.0 | 42.4 | 53.1 |
| **ONT_Zymo** | 243k | 84.0 | 71.2 | 77.0 | 65.9 | 80.3 | 66.5 | 88.2 | 88.0 | 88.1 | 83.7 | 42.5 | 49.8 |
| **PB_ATCC** | 95k | 96.8 | 93.5 | 99.6 | 99.0 | 100 | - | 100 | 100 | 100 | 100 | 72.6 | 77.0 |
| **PB_Zymo** | 117k | 98.0 | 97.0 | 97.0 | 75.2 | 91.3 | - | 98.4 | 92.8 | 96.1 | 89.5 | 81.1 | 58.1 |
| **Sample10** | 2.9M | 44.5 | 48.2 | 34.7 | 19.8 | 34.0 | - | 48.6 | 49.2 | 49.4 | 27.9 | 25.8 | 16.2 |
| **Sample20** | 3.2M | 28.4 | 31.4 | 17.1 | 14.3 | 12.2 | - | 23.6 | 25.3 | 25.4 | 8.7 | 11.0 | 4.6 |
| **Sample21** | 4.8M | 51.0 | 54.3 | 42.5 | 28.7 | 35.8 | - | 57.9 | 59.4 | 59.5 | 32.8 | 28.7 | 18.45 |
| **SRR15489009** | 956k | 84.5 | 85.6 | 81.6 | 51.0 | 59.5 | - | 87.9 | 77.4 | 84.1 | 53.7 | 87.3 | - |
| **SRR15489011** | 818k | 82.6 | 84.1 | 76.2 | 37.0 | 44.4 | - | 84.3 | 68.8 | 78.3 | 39.3 | 87.9 | 41.5 |
| **SRR15489017** | 1.1M | 81.9 | 82.7 | 79.8 | 44.0 | 55.2 | - | 86.9 | 75.2 | 82.7 | 48.6 | 86.2 | - |

### Supplementary Table 4

**The number of species detected for all real datasets and all tools.** For mock community datasets, the correct number of species is shown in parentheses after the dataset name.

| **Dataset**  **/ tool** | **Kraken2** | **Bracken** | **Centrifuge** | **CLARK** | **CLARK-S** | **Metamaps** | **MEGAN-N** | **deSAMBA** | **Minimap2 align** | **Minimap2 map** | **Ram** | **Kaiju** | **MEGAN -P** |
| --- | --- | --- | --- | --- | --- | --- | --- | --- | --- | --- | --- | --- | --- |
| **ONT_Zymo (10)** | 3224 | 114 | 3449 | 695 | 44 | 496 | 311 | 138 | 98 | 129 | 67 | 2737 | 863 |
| **PB_ATCC (20)** | 117 | 45 | 66 | 70 | 27 | 28 | - | 121 | 46 | 50 | 43 | 94 | 92 |
| **PB_Zymo (17)** | 530 | 102 | 794 | 225 | 50 | 72 | - | 195 | 105 | 132 | 97 | 379 | 185 |
| **Sample10** | 5487 | 4775 | 5538 | 2542 | 418 | 4973 | - | 1017 | 1051 | 1504 | 432 | 7102 | 3046 |
| **Sample20** | 5459 | 4560 | 5523 | 1736 | 452 | 4562 | - | 659 | 786 | 996 | 316 | 6868 | 2801 |
| **Sample21** | 5440 | 4944 | 5510 | 2220 | 375 | 4679 | - | 1010 | 999 | 1390 | 409 | 6837 | 3134 |
| **SRR15489009** | 4625 | 2011 | 5177 | 3785 | 294 | 415 | - | 2197 | 1904 | 4705 | 306 | 3184 | - |
| **SRR15489011** | 4599 | 2452 | 5193 | 3768 | 338 | 460 | - | 2049 | 1870 | 4663 | 351 | 3071 | 1145 |
| **SRR15489017** | 4784 | 2078 | 5260 | 3992 | 347 | 453 | - | 2383 | 2173 | 4855 | 345 | 3506 | - |

### Supplementary Table 5

**Composition of Zymo and ATCC standards.**

| **Zymo Microbial Community Standard (D6300) - ONT_Zymo dataset** | | **Zymo Gut Microbiome Standard (D6331) - PB_Zymo dataset** | | **ATCC MSA-1003 20 Strain Staggered Mix Genomic Material - PB_ATCC dataset** | |
| --- | --- | --- | --- | --- | --- |
| **Species** | **Theoretical Abun (%)** | **Species** | **Theoretical Abun (%)** | **Species** | **Theoretical Abun (%)** |
| **Pseudomonas aeruginosa** | **12** | **Faecalibacterium prausnitzii** | **14** | **Acinetobacter baumannii (ATCC 17978)** | **0.18** |
| **Escherichia coli** | **12** | **Veillonella rogosae** | **14** | **Bacillus pacificus (ATCC 10987)** | **1.80** |
| **Salmonella enterica** | **12** | **Roseburia hominis** | **14** | **Bacteroides vulgatus (ATCC 8482)** | **0.02** |
| **Lactobacillus fermentum** | **12** | **Bacteroides fragilis** | **14** | **Bifidobacterium adolescentis (ATCC 15703)** | **0.02** |
| **Enterococcus faecalis** | **12** | **Prevotella corporis** | **6** | **Clostridium beijerinckii (ATCC 35702)** | **1.80** |
| **Staphylococcus aureus** | **12** | **Bifidobacterium adolescentis** | **6** | **Cutibacterium acnes (ATCC 11828)** | **0.18** |
| **Listeria monocytogenes** | **12** | **Fusobacterium nucleatum** | **6** | **Deinococcus radiodurans (ATCC BAA-816)** | **0.02** |
| **Bacillus subtilis** | **12** | **Lactobacillus fermentum** | **6** | **Enterococcus faecalis (ATCC 47077)** | **0.02** |
| **Saccharomyces cerevisiae** | **2** | **Clostridioides difficile** | **1.5** | **Escherichia coli (ATCC 700926)** | **18.0** |
| **Cryptococcus neoformans** | **2** | **Akkermansia muciniphila** | **1.5** | **Helicobacter pylori (ATCC 700392)** | **0.18** |
|  | | **Methanobrevibacter smithii** | **0.1** | **Lactobacillus gasseri (ATCC 33323)** | **0.18** |
|  |  | **Salmonella enterica** | **0.01** | **Neisseria meningitidis (ATCC BAA-335)** | **0.18** |
|  |  | **Enterococcus faecalis** | **0.001** | **Porphyromonas gingivalis (ATCC 33277)** | **18.0** |
|  |  | **Clostridium perfringens** | **0.0001** | **Pseudomonas aeruginosa (ATCC 9027)** | **1.80** |
|  |  | **Escherichia coli (JM109)** | **2.8** | **Rhodobacter sphaeroides (ATCC 17029)** | **18.0** |
|  |  | **Escherichia coli (B-3008)** | **2.8** | **Schaalia odontolytica (ATCC 17982)** | **0.02** |
|  |  | **Escherichia coli (B-2207)** | **2.8** | **Staphylococcus aureus (ATCC BAA-1556)** | **1.80** |
|  |  | **Escherichia coli (B-766)** | **2.8** | **Staphylococcus epidermidis (ATCC 12228)** | **18.0** |
|  |  | **Escherichia coli (B-1109)** | **2.8** | **Streptococcus agalactiae (ATCC BAA-611)** | **1.80** |
|  |  | **Candida albicans** | **1.5** | **Streptococcus mutans (ATCC 700610)** | **18.0** |
|  |  | **Saccharomyces cerevisiae** | **1.4** |  | |

### Supplementary Table 6

**Genome lengths of species having more than 1% of reads classified for three real datasets.**

The table shows the average genome lengths of species in Mb that have more than 1% of reads classified to them by any tool and the average percentage of reads classified to those species.

| **ONT_Zymo** | | | **PB_ATCC** | | | **PB_Zymo** | | |
| --- | --- | --- | --- | --- | --- | --- | --- | --- |
| **tax id** | **genome length (Mb)** | **percentage of reads** | **tax id** | **genome length (Mb)** | **percentage of reads** | **tax id** | **genome length (Mb)** | **percentage of reads** |
| 1613 | 2.01 | 16.47 | 1063 | 4.62 | 31.48 | 39778 | 1.94 | 20.5 |
| 1639 | 2.97 | 16.32 | 1309 | 1.98 | 15.62 | 817 | 5.27 | 17.55 |
| 1351 | 2.97 | 15.11 | 562 | 5.13 | 14.93 | 562 | 5.13 | 10.5 |
| 1280 | 2.84 | 13.61 | 837 | 2.33 | 12.48 | 29466 | 2.13 | 8.04 |
| 28901 | 4.81 | 10.52 | 1282 | 2.51 | 10.99 | 851 | 2.42 | 5.47 |
| 562 | 5.13 | 9.37 | 287 | 6.6 | 2.38 | 853 | 2.8 | 5.18 |
| 287 | 6.6 | 7.66 | 1396 | 5.76 | 1.78 | 301301 | 3.06 | 4.63 |
| 1423 | 4.13 | 7.55 | 1311 | 2.08 | 0.51 | 28131 | 2.78 | 4.12 |
| 1963032 | 4.05 | 0.79 |  | | | 1613 | 2.01 | 2.35 |
| 72361 | 3.94 | 0.32 |  |  |  | 1680 | 2.2 | 2.19 |
| 1978467 | 6.22 | 0.14 |  |  |  | 239935 | 2.76 | 2.1 |
|  | | |  |  |  | 1496 | 4.15 | 1.86 |
|  |  |  |  |  |  | 39777 | 2.09 | 1.43 |
|  |  |  |  |  |  | 2682456 | 2.12 | 0.96 |
|  |  |  |  |  |  | 2682455 | 1.95 | 0.92 |
|  |  |  |  |  |  | 28133 | 2.69 | 0.19 |
|  |  |  |  |  |  | 28127 | 3.03 | 0.17 |

### Supplementary Table 7

**Isolate datasets used for testing dataset construction**. Data was downloaded from NCBI SRA (Sequence Read Archive), Sanger Institute pages (NCTC 3000) and PacBio GitHub DevNet pages (Microbial multiplexing dataset and Human Genome in a Bottle dataset). Isolates are identified by their identifier in the corresponding database, given below:

| **Nanopore isolate datasets** | | |
| --- | --- | --- |
| **Identifier** | **Source** | **Species** |
| SRR5891470 | NCBI | *Acinetobacter baumannii* |
| SRR6255797 | NCBI | *Amphiprion ocellaris* |
| DRR095709 | NCBI | *Apis cerana japonica* |
| ERR2173373 | ENA | *Arabidopsis thaliana* |
| SRR5817726 | NCBI | *Bacteroides caccae* |
| SRR5817721 | NCBI | *Blautia hansenii DSM 20583* |
| SRR5434252 | NCBI | *Borreliella burgdorferi* |
| SRR5457531 | NCBI | *Clostridioides difficile* |
| SRR5817725 | NCBI | *Enterocloster bolteae* |
| DRR048282 | NCBI | *Dengue virus* |
| DRR129651 | NCBI | *Vibrio litoralis DSM 17657* |
| DRR129656 | NCBI | *Vibrio aphrogenes* |
| DRR129658 | NCBI | *Vibrio algivorus* |
| DRR129660 | NCBI | *Vibrio casei* |
| DRR138509 | NCBI | *Homo sapiens* |
| DRR138514 | NCBI | *Homo sapiens* |
| DRR154114 | NCBI | *Streptococcus pneumoniae* |
| DRR155409 | NCBI | *Staphylococcus aureus* |
| DRR161061 | NCBI | *Staphylococcus argenteus* |
| DRR164908 | NCBI | *Leptotrichia trevisanii* |
| ERR1993378 | ENA | *Nippostrongylus brasiliensis* |
| ERR2014660 | EN | *Solanum americanum* |
| SRR6118140 | NCBI | *Escherichia coli* |
| SRR5890147 | NCBI | *Helcococcus kunzii* |
| SRR5902346 | NCBI | *Homo sapiens* |
| SRR6348587 | NCBI | *Klebsiella pneumoniae* |
| SRR5812844 | NCBI | *Lomentospora prolificans* |
| SRR5319868 | NCBI | *Maccullochella peelii* |
| SRR6192842 | NCBI | *Melanaphis sacchari* |
| SRR5997379 | NCBI | *Mesoplasma chauliocola* |
| SRR6129216 | NCBI | *Mesoplasma florum* |
| SRR6082028 | NCBI | *Mesoplasma lactucae ATCC 49193* |
| SRR5889392 | NCBI | *Oryza coarctata* |
| SRR5851902 | NCBI | *Rhodotorula mucilaginosa* |
| SRR6059708 | ENA | *Saccharomyces cerevisiae* |
| ERR2177070 | ENA | *Solanum pennellii* |
| **PacBio isolate datasets** | | |
| **Identifier** | **Source** | **Species** |
| NCTC10116 | NCTC | *Acholeplasma laidlawii* |
| NCTC12156 | NCTC | *Acinetobacter baumannii* |
| NCTC10308 | NCTC | *Acinetobacter johnsonii* |
| NCTC10221 | NCTC | *Actinobacillus ureae* |
| NCTC8251 | NCTC | *Aerococcus viridans* |
| NCTC8049 | NCTC | *Aeromonas hydrophila* |
| NCTC5906 | NCTC | *Aggregatibacter aphrophilus ATCC 33389* |
| Arabidopsis P5C3 | PacBio GitHub DevNet | *Arabidopsis thaliana* |
| ACTC 14579 | PacBio GitHub DevNet | *Bacilus cereus 971* |
| Bacillus subtilis W23 | PacBio GitHub DevNet | *Bacillus subtilis* |
| NCTC12860 | PacBio | *Bartonella grahamii* |
| NCTC11829 | PacBio | *Blautia producta* |
| NCTC2611 | PacBio | *Brevibacillus brevis* |
| ATCC 25416 | PacBio GitHub DevNet | *Burkholderia cepacia* |
| NCTC11168 | NCTC | *Campylobacter jejuni* |
| NCTC12120 | NCTC | *Cedecea neteri* |
| NCTC13790 | NCTC | *Citrobacter braakii* |
| NCTC11811 | NCTC | *Clostridium indolis* |
| SRX499318 | ENA | *Drosophila melanogaster* |
| NCTC11813 | NCTC | *Eggerthella lenta* |
| NCTC418 | NCTC | *Enterobacter aerogenes* |
| OG1RF ATCC 47077D-5 | PacBio GitHub DevNet | *Enterococcus faecalis* |
| escherichia coli K12 | PacBio GitHub DevNet | *Escherichia coli* |
| NCTC4450 | NCTC | *Escherichia coli* |
| ATCC 9637 | PacBio GitHub DevNet | *Escherichia coli W* |
| NCTC11401 | NCTC | *Fluoribacter gormanii* |
| NCTC10562 | NCTC | *Fusobacterium nucleatum* |
| NCTC11413 | NCTC | *Gallibacterium anatis* |
| NCTC10667 | NCTC | *Gordonia bronchialis* |
| NCTC11483 | NCTC | *Haemophilus ducreyi* |
| NCTC11931 | NCTC | *Haemophilus influenzae* |
| ATCC 700824 | PacBio GitHub DevNet | *Helicobacter pylori J99* |
| HG002 | PacBio GitHub DevNet | *Homo sapiens* |
| NCTC10816 | NCTC | *Jonesia denitrificans* |
| ATCC BAA-2146 | PacBio GitHub DevNet | *Klebsiella pneumoniae* |
| INF274 rs 2 | NCTC | *Klebsiella pneumoniae* |
| NCTC11040 | NCTC | *Kytococcus sedentarius* |
| NCTC13720 | NCTC | *Lactobacillus acidophillus* |
| NCTC13768 | NCTC | *Lactobacillus brevis* |
| NCTC11417 | NCTC | *Legionella pneumophila* |
| NCTC11994 | NCTC | *Listeria monocytogenes* |
| ATCC 43576 | PacBio GitHub DevNet | *Methanocorpusculum labreanum Z* |
| NCTC2665 | NCTC | *Micrococcus luteus* |
| NCTC11656 | NCTC | *Mobiluncus curtisii* |
| NCTC10437 | NCTC | *Mycobacterium aurum* |
| ATCC 700532 | PacBio GitHub DevNet | *Neisseria meningitidis FAM18* |
| NCTC11134 | NCTC | *Nocardia farcinica* |
| NCTC12168 | NCTC | *Ochrobactrum anthropi* |
| NCTC10343 | NCTC | *Paenibacillus polymyxa* |
| NCTC9381 | NCTC | *Pantoea agglomerans* |
| NCTC10204 | NCTC | *Pasteurella multocida* |
| NCTC11647 | NCTC | *Photobacterium damselae* |
| SRX3923117 | NCBI | *Pseudomonas koreensis P19E3* |
| rhodopseudomonas palustris | PacBio GitHub DevNet | *Rhodopseudomonas palustris* |
| NCTC10917 | NCTC | *Rothia dentocariosa* |
| NCTC12419 | NCTC | *Salmonella bongori* |
| NCTC10384 | NCTC | *Salmonella enterica* |
| NCTC12961 | NCTC | *Serratia plymuthica* |
| NCTC13711 | NCTC | *Staphylococcus argenteus* |
| NCTC13277 | NCTC | *Staphylococcus aureus* |
| ATCC 25923 | PacBio GitHub DevNet | *Staphylococcus aureus subsp aureus* |
| NCTC7466 | NCTC | *Streptococcus pneumoniae* |
| ATCC 35405 | PacBio GitHub DevNet | *Treponema denticola A* |
| NCTC11317 | NCTC | *Vibrio campbellii* |
| NCTC10938 | NCTC | *Yersinia enterocolitica* |
| NCTC12986 | NCTC | *Yersinia ruckeri* |

### Supplementary Table 8

**Tie handling while processing mappers output.** Read-level classification for mappers was obtained from PAF and SAM files, using a single classification for each read. In the case of ties, the first alignment was used. The table shows some relevant statistics for reads that have more than one equally scored primary alignment.

| Dataset | Tool | Percentage of reads that have more than one equally scored primary alignment | Percentage of reads with more than one equally scored primary alignment that was correctly classified | Percentage of reads that have more than one equally scored primary alignment that would have been correctly classified if another primary alignment was chosen |
| --- | --- | --- | --- | --- |
| **ONT1** | **Minimap2** | 2.29% | 10.06% | 82.06% |
|  | **Ram** | 2.05% | 5.43% | 90.86% |
| **ONT2** | **Minimap2** | 0.40% | 32.11% | 16.72% |
|  | **Ram** | 0.25% | 45.75% | 21.86% |
| **PB1** | **Minimap2** | 0.83% | 15.57% | 70.07% |
|  | **Ram** | 0.70% | 33.00% | 55.29% |
| **PB2** | **Minimap2** | 1.38% | 2.90% | 21.21% |
|  | **Ram** | 0.28% | 36.73% | 50.91% |
| **PB3** | **Minimap2** | 0.03% | 22.22% | 11.11% |
|  | **Ram** | 0.02% | 56.25% | 18.75% |
| **PB4** | **Minimap2** | 0.91% | 12.31% | 48.46% |
|  | **Ram** | 0.84% | 28.79% | 31.75% |

### Supplementary Table 9

**Abundance estimation error for low abundance species.** The table shows the abundance estimation error calculated as the L1 distance between the expected value and the value reported by tools. The error is calculated only for species for which the expected value for abundance is 1% or less. The table also shows the number of correctly detected low-abundance species (in parentheses).

| **Dataset**  **/ tool** | **Total abundance and number of low abundance species** | **Kraken2** | **Bracken** | **Centrifuge** | **CLARK** | **CLARK-S** | **Metamaps** | **MEGAN-N** | **deSAMBA** | **Minimap2 align** | **Minimap2 map** | **Ram** | **Kaiju** | **MEGAN-P** |
| --- | --- | --- | --- | --- | --- | --- | --- | --- | --- | --- | --- | --- | --- | --- |
| **ONT1** | 2.1  (7) | 1.40  **(7)** | 2.81  (6) | 1.29  **(7)** | 0.56  **(7)** | 0.93  **(7)** | 0.62  **(7)** | 1.47  **(7)** | 0.67  **(7)** | 1.17  **(7)** | 1.13  **(7)** | 1.20  **(7)** | 0.61  **(7)** | **0.21**  **(7)** |
| **PB4** | 10.0  (27) | 2.70  **(25)** | 5.92  (23) | 2.80  (24) | 2.84  **(25)** | 3.90  (23) | 3.08  (23) | 2.54  (24) | 2.53  (24) | **2.38**  (24) | 2.41  **(25)** | 2.63  (24) | 5.07  (24) | 3.4  (24) |
| **PB_ATCC** | 1.0  (10) | 0.61  **(10)** | **0.51**  (7) | 0.64  **(10)** | 0.60  **(10)** | 0.61  **(10)** | 0.60  **(10)** | - | 0.62  **(10)** | 0.60  **(10)** | 0.60  **(10)** | 0.60  **(10)** | 0.72  **(10)** | 0.68  **(10)** |
| **PB_Zymo** | 0.8  (4) | 0.039  **(4)** | 0.026  (2) | 0.019  (3) | 0.014  (3) | 0.088  (3) | 0.009  (3) | - | 0.037  (3) | **0.008**  (3**)** | 0.015  (3) | 0.013  (3) | 0.033  (3) | 0.175  (3) |

### Supplementary Table 10

**List of species in the datasets that are not in the database.** The table lists all the species in datasets that are not found in the nucleotide or protein database and their true abundances in corresponding datasets. Only synthetic and mock datasets are shown in the table since, for them, the species that are present in the dataset are known.

| **dataset** | **tax id** | **species name** | **true abundance** | **missing in NT database** | **missing in NR database** |
| --- | --- | --- | --- | --- | --- |
| ONT1 | 1667024 | Vibrio algivorus | 10.82% | YES | YES |
|  | 335972 | Vibrio litoralis | 7.33% | YES | YES |
|  | 40091 | Helcococcus kunzii | 6.77% | YES | **NO** |
| ONT2 | 1667024 | Vibrio algivorus | 12.31% | YES | YES |
|  | 335972 | Vibrio litoralis | 10.62% | YES | YES |
| PB2 | 7227 | Drosophila melanogaster | 1.73% | YES | YES |
| PB4 | 69825 | Lacrimispora indolis | 2.03% | YES | YES |
|  | 723 | Actinobacillus ureae | 1.34% | YES | YES |
|  | 464 | Fluoribacter gormanii | 0.11% | YES | YES |
| PB_Zymo | 423477 | Veillonella rogosae | 19.94% | YES | YES |
|  | 28128 | Prevotella corporis | 6.26% | YES | YES |
|  | 4932 | Saccharomyces cerevisiae | 0.32% | YES | YES |
|  | 5476 | Candida albicans | 0.31% | YES | YES |
| ONT_Zymo | 4932 | Saccharomyces cerevisiae | 0.57% | YES | YES |
|  | 5207 | Cryptococcus neoformans | 0.37% | YES | YES |
| SRR15489009 | 2109690 | Lachnospiraceae bacterium Choco86 | - | NO | YES |
|  | 2109691 | Lachnospiraceae bacterium GAM79 | - | NO | YES |
|  | 1898203 | Lachnospiraceae bacterium | - | NO | YES |
|  | 2594789 | Lachnospiraceae bacterium KGMB03038 | - | NO | YES |
|  | 2093742 | Lachnospiraceae bacterium KM106-2 | - | NO | YES |
|  | 712991 | Lachnospiraceae bacterium oral taxon 500 | - | NO | YES |
|  | 165179 | Prevotella copri | - | YES | NO |
|  | 33039 | Ruminococcus torques | - | YES | NO |
|  | 40520 | Blautia obeum | - | YES | NO |
|  | 1737424 | Blautia massiliensis | - | YES | NO |
|  | 410072 | Coprococcus comes | - | YES | NO |
|  | 39491 | Eubacterium rectale | - | NO | YES |

### Supplementary Table 11

**Abundance results for SRR15489009(PacBio) dataset.** The table lists abundances for the first 10 most abundant species for every tool for the SRR15489009 dataset sorted by name. MEGAN-N tool did not produce results for SRR15489009 and is omitted from this table. Cells with a higher discordance among tools are shaded in grey.

| **Species name** | **Kraken 2** | **Bracken** | **Centrifuge** | **CLARK** | **CLARK-S** | **Metamaps** | **DeSamba** | **Minimap2 align** | **Minimap2 map** | **ram** | **kaiju** | **MEGAN-P** |
| --- | --- | --- | --- | --- | --- | --- | --- | --- | --- | --- | --- | --- |
| Anaerobutyricum hallii | 3.04 | 2.55 | 3.16 | 3.67 | 4.07 | 3.43 | 2.55 | 3.32 | 2.91 | 3.03 | 2.38 | 2.76 |
| Anaerostipes hadrus | 4.61 | 5.02 | 3.91 | 5.32 | 8.18 | 7.11 | 4.45 | 6.49 | 5.15 | 7.33 | 4.08 | 6.19 |
| Bifidobacterium adolescentis | 3.03 | 2.56 | 2.98 | 3.18 | 6.23 | 5.37 | 2.80 | 4.86 | 3.49 | 5.73 | 2.73 | 5.22 |
| Bifidobacterium angulatum | 2.07 | 1.81 | 2.04 | 2.17 | 4.31 | 3.73 | 1.95 | 3.39 | 2.43 | 4.00 | 1.88 | 3.75 |
| Bifidobacterium longum | 1.00 | 0.84 | 0.96 | 1.04 | 1.97 | 1.71 | 0.93 | 1.56 | 1.14 | 1.85 | 0.94 | 1.73 |
| Blautia argi | 1.19 | 1.23 | 0.99 | 1.14 | 0.28 | 0.46 | 1.00 | 0.65 | 0.86 | 0.64 | 0.56 | 0.09 |
| Blautia massiliensis | 0.00 | 0.00 | 0.00 | 0.00 | 0.00 | 0.00 | 0.00 | 0.00 | 0.00 | 0.00 | 2.43 | 1.37 |
| Blautia obeum | 0.00 | 0.00 | 0.00 | 0.00 | 0.00 | 0.00 | 0.00 | 0.00 | 0.00 | 0.00 | 3.53 | 2.06 |
| Blautia producta | 0.63 | 0.53 | 0.65 | 0.63 | 0.06 | 0.07 | 0.63 | 0.24 | 0.53 | 0.15 | 0.27 | 0.10 |
| Blautia sp. SC05B48 | 11.79 | 9.39 | 9.56 | 11.13 | 13.60 | 10.16 | 10.95 | 10.42 | 11.42 | 7.96 | 1.61 | 0.73 |
| Collinsella aerofaciens | 2.57 | 2.20 | 2.51 | 2.74 | 5.32 | 4.63 | 2.42 | 4.10 | 2.99 | 4.87 | 2.33 | 4.48 |
| Coprococcus comes | 0.00 | 0.00 | 0.00 | 0.00 | 0.00 | 0.00 | 0.00 | 0.00 | 0.00 | 0.00 | 3.01 | 2.37 |
| Eubacterium  rectale | 0.00 | 0.00 | 0.00 | 0.00 | 0.00 | 6.10 | 2.21 | 5.84 | 4.99 | 5.96 | 0.00 | 0.00 |
| Faecalibacterium prausnitzii | 12.40 | 14.89 | 11.86 | 13.26 | 24.73 | 18.54 | 11.80 | 17.12 | 14.09 | 17.64 | 11.47 | 19.27 |
| Lachnospiraceae bacterium | 0.99 | 5.97 | 0.81 | 0.00 | 0.00 | 0.44 | 4.06 | 1.01 | 1.47 | 0.76 | 0.00 | 0.00 |
| Lachnospiraceae bacterium Choco86 | 3.84 | 3.13 | 2.52 | 0.00 | 0.00 | 1.98 | 2.82 | 2.16 | 2.58 | 1.75 | 0.00 | 0.00 |
| Lachnospiraceae bacterium GAM79 | 3.84 | 3.65 | 3.08 | 0.00 | 0.00 | 4.01 | 2.72 | 3.58 | 3.04 | 3.95 | 0.00 | 0.00 |
| Lachnospiraceae bacterium KGMB03038 | 0.48 | 0.45 | 0.33 | 0.00 | 0.00 | 0.01 | 0.29 | 0.02 | 0.22 | 0.02 | 0.00 | 0.00 |
| Lachnospiraceae bacterium KM106-2 | 0.43 | 0.37 | 0.36 | 0.00 | 0.00 | 0.20 | 0.22 | 0.22 | 0.21 | 0.32 | 0.00 | 0.00 |
| Lachnospiraceae bacterium oral taxon 500 | 0.08 | 0.07 | 0.04 | 0.00 | 0.00 | 0.00 | 0.02 | 0.02 | 0.02 | 0.02 | 0.00 | 0.00 |
| Prevotella copri | 0.00 | 0.00 | 0.00 | 0.00 | 0.00 | 0.00 | 0.00 | 0.00 | 0.00 | 0.00 | 8.32 | 12.85 |
| Roseburia hominis | 2.50 | 2.39 | 1.50 | 2.85 | 1.61 | 1.73 | 3.02 | 2.17 | 2.63 | 2.12 | 3.36 | 2.31 |
| Roseburia intestinalis | 5.52 | 5.03 | 5.42 | 6.63 | 7.43 | 7.24 | 6.30 | 6.34 | 6.35 | 6.44 | 6.06 | 7.36 |
| Ruminococcus  gnavus | 2.06 | 1.73 | 0.95 | 2.08 | 0.97 | 0.92 | 2.56 | 1.48 | 2.12 | 1.05 | 1.10 | 0.42 |
| Ruminococcus  torques | 0.00 | 0.00 | 0.00 | 0.00 | 0.00 | 0.00 | 0.00 | 0.00 | 0.00 | 0.00 | 3.72 | 4.07 |
| Streptococcus salivarius | 1.31 | 1.27 | 1.03 | 1.40 | 2.48 | 2.04 | 1.02 | 1.76 | 1.27 | 2.06 | 0.96 | 0.60 |
